# Pathogenic variants of sphingomyelin synthase SMS2 disrupt lipid landscapes in the secretory pathway

**DOI:** 10.1101/2022.04.02.486804

**Authors:** Tolulope Sokoya, Jan Parolek, Mads Møller Foged, Dmytro I. Danylchuk, Manuel Bozan, Bingshati Sarkar, Angelika Hilderink, Michael Philippi, Lorenzo D. Botto, Paulien A. Terhal, Outi Mäkitie, Jacob Piehler, Yeongho Kim, Christopher G. Burd, Andrey S. Klymchenko, Kenji Maeda, Joost C. M. Holthuis

## Abstract

Sphingomyelin is a dominant sphingolipid in mammalian cells. Its production in the *trans*-Golgi traps cholesterol synthesized in the ER to promote formation of a sphingomyelin/sterol gradient along the secretory pathway. This gradient marks a fundamental transition in physical membrane properties that help specify organelle identify and function. We previously identified mutations in sphingomyelin synthase SMS2 that cause osteoporosis and skeletal dysplasia. Here we show that SMS2 variants linked to the most severe bone phenotypes retain full enzymatic activity but fail to leave the ER owing to a defective autonomous ER export signal. Cells harboring pathogenic SMS2 variants accumulate sphingomyelin in the ER and display a disrupted transbilayer sphingomyelin asymmetry. These aberrant sphingomyelin distributions also occur in patient-derived fibroblasts and are accompanied by imbalances in cholesterol organization, glycerophospholipid profiles and lipid order in the secretory pathway. We postulate that pathogenic SMS2 variants undermine the capacity of osteogenic cells to uphold nonrandom lipid distributions that are critical for their bone forming activity.

## INTRODUCTION

Eukaryotic membranes consist of complex lipid mixtures, with amounts and ratios of the individual lipids showing marked variations between organelles and membrane leaflets (van Meer et al., 2008; Harayama and Riezman, 2018). Whereas some rare lipids contribute to organelle function by allowing stereospecific recognition through lipid binding proteins (Di Paolo and De Camilli, 2006), numerous recognition processes on or within organellar bilayers are determined by biophysical membrane properties that result from the collective behavior of the bulk lipids. Particularly striking are the lipid-induced changes in bilayer-thickness, lipid packing density and surface charge that accompany the transition from early to late organelles in the secretory pathway (Sharpe et al., 2010; Bigay and Antonny, 2012; Holthuis and Menon, 2014). These changes are highly conserved and provide specific cues for membrane proteins that govern vital processes such as protein secretion and signaling (Bigay and Antonny, 2012; Magdeleine et al., 2016; Zhou and Hancock, 2018). To defend the unique lipid mixtures of secretory organelles against erosion by vesicular transport, cells exploit cytosolic transfer proteins that enable specific lipids to bypass vesicular connections by mediating their monomeric exchange at contact sites between distinct organelles (Wong et al., 2019). Moreover, organelles like the ER harbor membrane property sensors that provide feedback to the lipid metabolic network to preserve their characteristic lipid composition when exposed to stress or metabolic insults (Radanović et al., 2018; Levental et al., 2020).

Sterols and sphingomyelin (SM) are prime examples of bulk membrane lipids that are unevenly distributed between secretory organelles (van Meer et al., 2008). Sterols are rare in the ER but abundant in the *trans*-Golgi and plasma membrane (PM). The bulk of SM is synthesized in the lumen of the *trans*-Golgi from ceramides supplied by the ER and delivered by vesicular transport to the PM, where it accumulates in the exoplasmic leaflet (Hanada et al., 2003). SM is the preferred binding partner of cholesterol (Slotte, 2013). About one third of the total cholesterol pool in the PM is sequestered by SM (Das et al., 2014; Endapally et al., 2019). Besides influencing cellular cholesterol homeostasis, SM contributes to an enhanced packing density and thickening of *trans*-Golgi and PM bilayers. This, in turn, may modulate protein sorting by hydrophobic mismatching of membrane spans (Munro, 1995; Quiroga et al., 2013). Moreover, an asymmetric distribution of SM across late secretory and endolysosomal bilayers is relevant for an optimal repair of damaged organelles. Lysosome wounding by chemicals or bacterial toxins triggers a rapid Ca^2+^-activated scrambling and cytosolic exposure of SM (Ellison et al., 2020; Niekamp et al., 2022). Subsequent conversion of SM to ceramides by neutral SMases on the cytosolic surface of injured lysosomes promotes their repair, presumably by driving an inverse budding of the damaged membrane area in a process akin to ESCRT-mediated formation of intraluminal vesicles. This SM-based membrane restoration pathway functions independently of ESCRT and may also operate at the PM (Niekamp et al., 2022).

SM biosynthesis in mammals is mediated by SM synthase 1 (SMS1) and SMS2. Both enzymes act as phosphatidylcholine (PC):ceramide phosphocholine transferases, which catalyze the transfer of the phosphorylcholine head group from PC onto ceramide to generate SM and diacylglycerol (DAG) (Huitema et al., 2004). SMS1 resides in the *trans*-Golgi, and its deficiency in mice causes mitochondrial dysfunction and disrupts insulin secretion (Yano et al., 2011, 2013). SMS2 resides both in the *trans*-Golgi and at the PM. Its deficiency ameliorates diet-induced obesity and insulin resistance (Li et al., 2011; Mitsutake et al., 2011; Sugimoto et al., 2016; Kim et al., 2018). Removal of SMS1 or SMS2 has only a minor impact on ceramide, DAG and SM pools in tissues or cells, and the mechanisms underlying the phenotypes observed in SMS1 and SMS2 knockout mice are not well understood. Besides SMS1 and SMS2, mammalian cells contain an ER-resident and SMS-related protein (SMSr), which displays phospholipase C activity and synthesizes trace amounts of the SM analog ceramide phosphoethanolamine (Vacaru et al., 2009; Murakami and Sakane, 2021).

We previously reported that SMS2 is highly expressed in bone and identified heterozygous mutations in the SMS2-encoding gene (*SGMS2*) as the underlying cause of a clinically described autosomal dominant genetic disorder – osteoporosis with calvarial doughnut lesions (OP-CDL: OMIM #126550) (Pekkinen et al., 2019). The clinical presentations of OP-CDL range from childhood-onset osteoporosis with low bone mineral density, skeletal fragility and sclerotic doughnut-shaped lesions in the skull to a severe spondylometaphyseal dysplasia with neonatal fractures, long-bone deformities, and short stature. The milder phenotype is associated with the nonsense variant p.Arg50*, which gives rise to a truncated but catalytically active enzyme that mislocalizes to the *cis/medial*-Golgi (T. Sokoya and J.C.M. Holthuis, unpublished). However, the most severe phenotypes are associated with two closely localized missense variants, p.Ile62Ser and p.Met64Arg. Interestingly, these missense variants enhance *de novo* SM biosynthesis by blocking ER export of enzymatically active SMS2 (Pekkinen et al., 2019). This suggests that OP-CDL in patients with pathogenic SMS2 variants is not due to a reduced capacity to synthesize SM but rather a consequence of mistargeting bulk SM production to an early secretory organelle. How this affects the contrasting lipid landscapes and membrane properties in the secretory pathway remains to be established.

In this work, we used genetically engineered cell lines and OP-CDL patient-derived fibroblasts to address the impact of pathogenic SMS2 variants p.Ile62Ser and p.Met64Arg on the lipid composition, transbilayer arrangement, and packing density of early and late secretory organelles. Toward this end, we combined shotgun lipidomics on purified organelles with the application of lipid biosensors and targeted solvatochromic fluorescent probes in live cells. We show that cells harbouring pathogenic SMS2 variants accumulate PM-like amounts of SM in the ER and display a disrupted transbilayer SM asymmetry. This is accompanied by significant imbalances in cholesterol organization and membrane lipid order. We also find that pathogenic SMS2 variants cause marked changes in the ER glycerophospholipid profile, including an enhanced phospholipid desaturation and rise in cone-shaped ethanolamine-containing phospholipids, potentially reflecting an adaptive cellular response to counteract SM-mediated rigidification of the ER bilayer. Our data indicate that pathogenic SMS2 variants profoundly undermine the cellular capacity to uphold nonrandom lipid distributions in the secretory pathway that may be critical for the bone forming activity of osteogenic cells.

## RESULTS

### The IXMP motif in SMS2 is part of an autonomous ER export signal

The most severe clinical presentations of OP-CDL are associated with the SMS2 missense variants p.I62S and p.M64R, which cause retention of a functional enzyme in the ER (Pekkinen et al., 2019). Ile62 and Met64 are part of a conserved sequence motif, IXMP, which is located 13-14 residues upstream of the first membrane span in both SMS1 and SMS2 (**Fig. 1a, b**). We reasoned that this motif may be part of an ER export signal, which could explain its absence in the ER-resident SMS family member SMSr (Vacaru et al., 2009). To test this idea, we generated FLAG-tagged SMS2 constructs in which Ile62 or Met64 was replaced with Ser or Arg, respectively. Upon their transfection in HeLa cells, the subcellular distribution of the SMS2 variants was determined by fluorescence microscopy using antibodies against the FLAG tag and ER-resident protein calnexin. In agreement with our previous findings (Pekkinen et al., 2019), SMS2^I62S^ and SMS2^M64R^ were each retained in the ER, in contrast to wildtype SMS2, which localized to the Golgi and PM (**Fig. 1c**). We then asked whether the IXMP motif in SMS2 can mediate ER export independently of other sorting information. To address this, we created a FLAG-tagged chimera protein, SMSr-SMS2_11-77_, in which the region linking the *N*-terminal SAM domain and first membrane span of SMSr was replaced with the IXMP-containing cytosolic tail of SMS2 (**Fig. 1a**). Contrary to SMSr, SMSr-SMS211-77 localized to the Golgi. However, SMSr-SMS2_11-77_ variants in which Ile62 or Met64 was replaced with Ser or Arg, respectively, were retained in the ER (**Fig. 1d**). This indicates that the IXMP motif in SMS2 is part of an autonomous ER export signal.

**Fig. 1.**
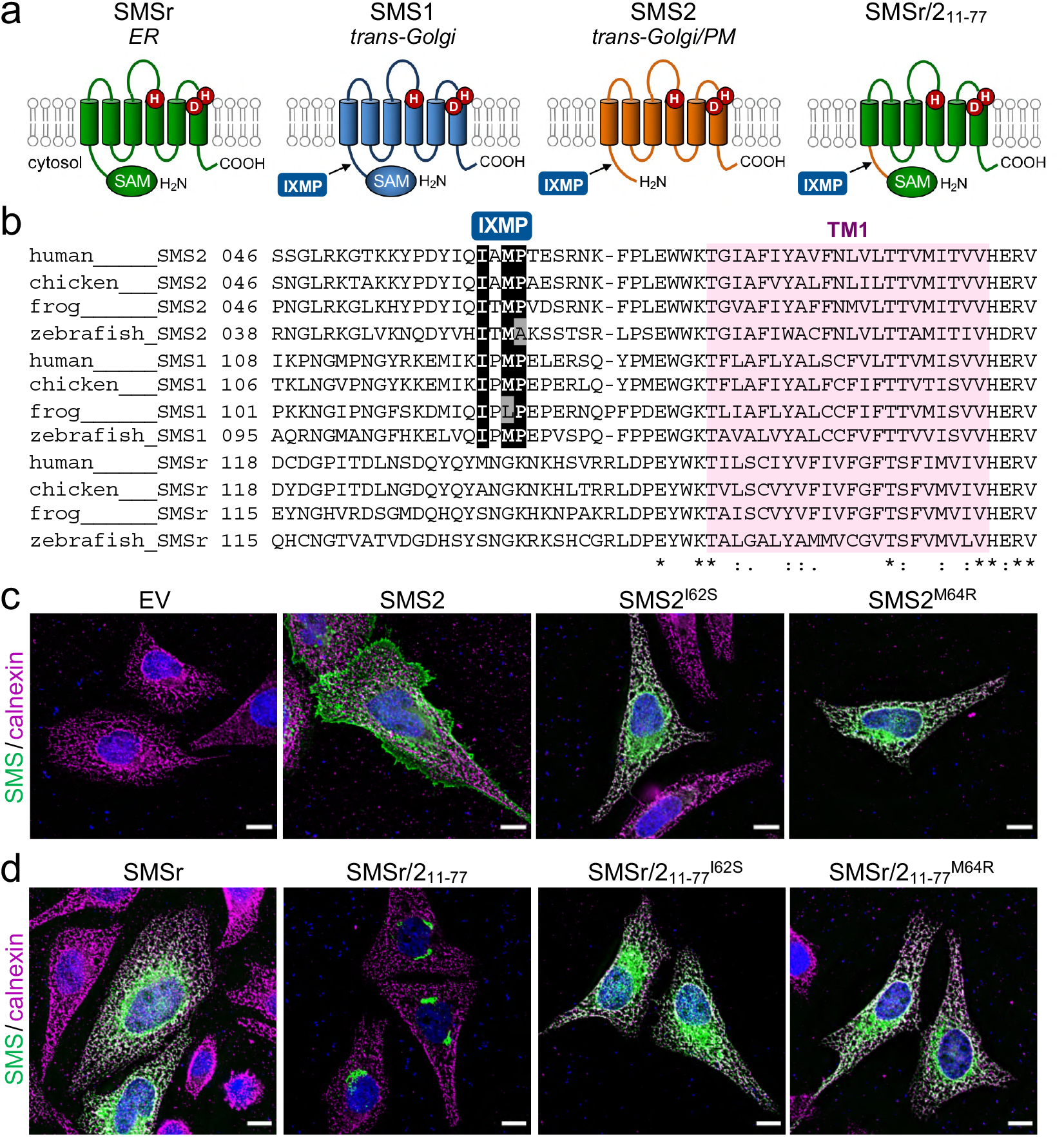
SMS2 contains an autonomous ER export signal. (**a**) Membrane topology of SMS family members and chimeric protein SMSr/2_11-77_. Active site residues are marked in red. The position of a conserved IXMP sequence motif is marked by an arrow. (**b**) Sequence alignment of the region immediately upstream of the first membrane span (TM1) in vertebrate SMS family members. Note that human SMS2 residues Ile62 and Met64 are part of the IXMP sequence motif, which is conserved in SMS1 and SMS2, but not SMSr, across different vertebrate species. (**c**) HeLa cells transfected with empty vector (EV) or FLAG-tagged SMS2, SMS2^I62S^ or SMS2^M64R^ were fixed, immunostained with α-FLAG (*green*) and a-calnexin (*magenta*) antibodies, counterstained with DAPI (*blue*) and imaged by DeltaVision microscopy. (**d**) HeLa cells transfected with FLAG-tagged SMSr, SMSr/2_11-77_, SMSr/2_11-77_^I62S^ or SMSr/2n-77^M64R^ were fixed, immunostained with α-FLAG (*green*) and α-calnexin (*magenta*) antibodies, counterstained with DAPI (*blue*) and imaged by DeltaVision microscopy. Scale bar, 10 μm.

### Pathogenic SMS2 variants mediate bulk production of SM in the ER

Metabolic labeling of patient-derived fibroblasts with ^14^C-choline showed that missense SMS2 variants p.I62S and p.M64R cause a marked increase in the rate of *de novo* SM biosynthesis (Pekkinen et al., 2019). To directly test the impact of these pathogenic mutations on the biosynthetic capacity of SMS2, we stably transduced SMS1/2 double knockout (ΔSMS1/2) HeLa cells with doxycycline-inducible expression constructs encoding FLAG-tagged SMS2, SMS2^I62S^, SMS2^M64R^ or their enzyme dead isoforms SMS2^D276A^, SMS2^I62S/D276A^ and SMS2^M64R/D276A^, respectively. After treatment of cells with doxycycline for 16 h, SMS2 expression was verified by immunoblot analysis and fluorescence microscopy (**Fig. 2a** and **b; Fig. S1**). Next, control and doxycycline-treated cells were metabolically labelled with a clickable sphingosine analogue for 16 h, subjected to total lipid extraction, click reacted with the fluorogenic dye 3-azido-7-hydroxycoumarin, and analyzed by TLC. This revealed that doxycycline-induced expression of SMS2^I62S^ and SMS2^M64R^, but not their enzyme-dead isoforms, fully restored SM biosynthesis in ΔSMS1/2 cells (**Fig. 2c**).

**Fig. 2.**
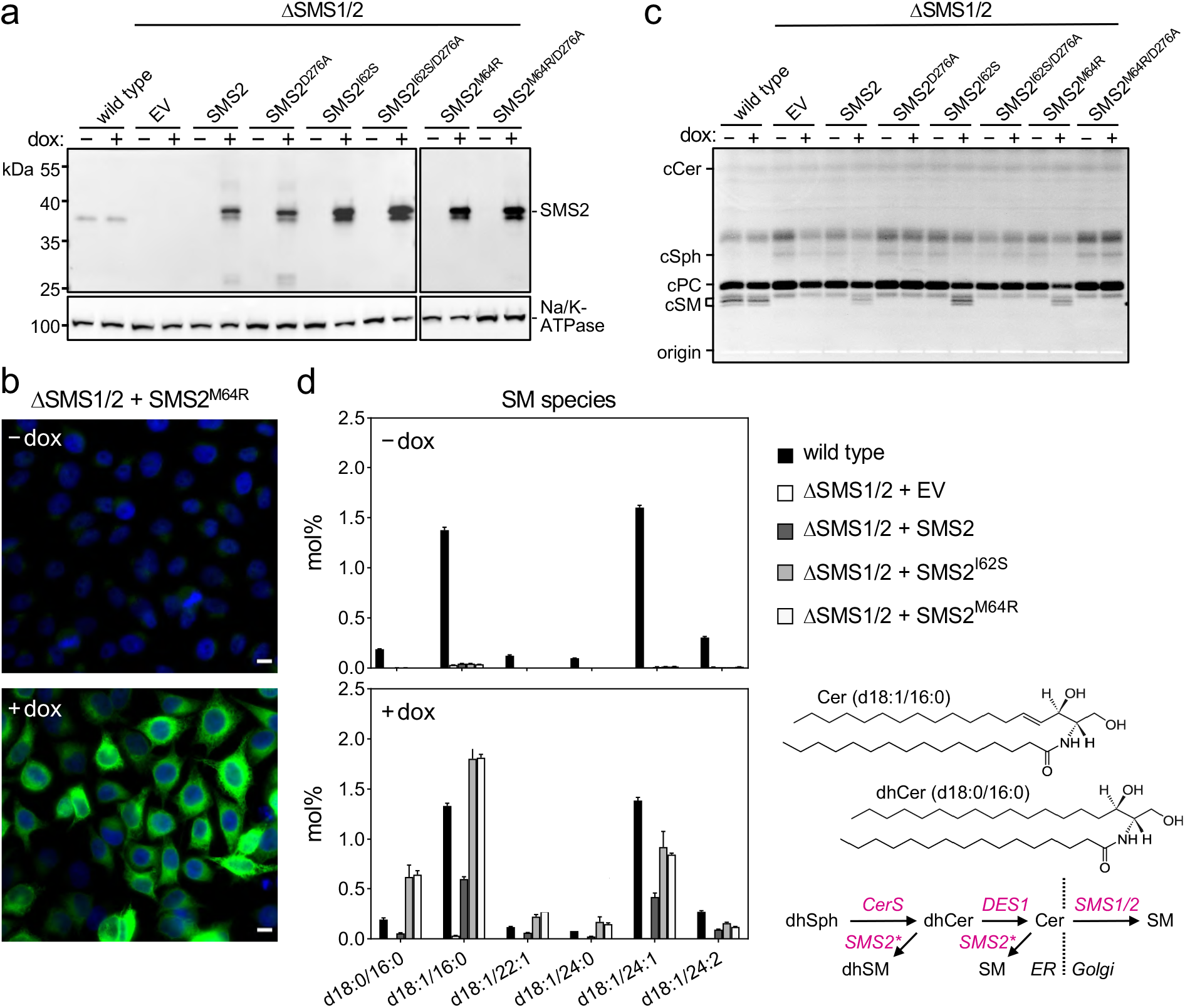
Pathogenic SMS2 variants support bulk production of SM in the ER. (**a**) HeLa SMS1/2 double KO (ΔSMS1/2) cells transduced with doxycycline-inducible constructs encoding FLAG-tagged SMS2, SMS2^I62S^, SMS2^M64R^ or their enzyme-dead isoforms (D276A) were grown for 16 h in the absence or presence of 1 μg/ml doxycycline and then subjected to immunoblot analysis using α-SMS2 and α-Na/K-ATPase antibodies. Wildtype HeLa cells served as control. (**b**) ΔSMS1/2 cells transduced with doxycycline-inducible FLAG-tagged SMS2^M64R^ were treated as in (a), fixed, immunostained with a-FLAG antibody (*green*), counterstained with DAPI (*blue*) and imaged by conventional fluorescence microscopy. Scale bar, 10 μm. (**c**) Cells treated as in (a) were metabolically labelled with a clickable sphingosine analogue for 16 h, subjected to total lipid extraction, click reacted with the fluorogenic dye 3-azido-7-hydroxycoumarin and analyzed by TLC. (**d**) SM species in total lipid extracts of cells treated as in (a) were quantified by LC-MS/MS and expressed as mol% of total phospholipid analyzed. Note that the rise in dihydroSM (dhSM, d18:0/16:0) in ΔSMS1/2 cells expressing SMS2^I62S^ or SMS2^M64R^ (SMS2*) is likely due to competition between ER-resident ceramide desaturase (DES1) and SMS2* for dihydroceramide (dhCer, d18:0/16:0), which is synthesized *de novo* by ceramide synthase (CerS) from dihydrosphingosine (dhSph).

Quantitative mass spectrometric analysis of total lipid extracts from wildtype and ΔSMS1/2 cells revealed that removal of SMS1 and SMS2 wiped out the entire cellular SM pool and caused a four-fold increase in glycosphingolipid (GSL) levels, consistent with a competition between Golgi-resident SM and glucosylceramide (GlcCer) synthases for ceramide substrate (**Fig. 2d**; **Fig. S2**). In ΔSMS1/2 cells transduced with pathogenic SMS2^I62S^ or SMS2^M64R^, addition of doxycycline fully restored the SM pool. This was accompanied by a decrease in GSL levels. Doxycycline-induced expression of SMS2 only partially restored the SM pool, presumably because SMS2, unlike its pathogenic isoforms, has no direct access to ER-derived ceramides and must compete with GlcCer synthase for ceramides delivered to the Golgi. Moreover, ΔSMS1/2 cells expressing SMS2^I62S^ or SMS2^M64^ contained 3- to 4-fold higher levels of dihydroceramide (Cer d18:0/16:0) and dihydroceramide-based SM (SM d18:0/16:0) than wildtype or SMS2-expressing ΔSMS1/2 cells (**Fig. 2d; Fig. S2b**), which suggests that ER-resident pathogenic SMS2 variants compete with ceramide desaturase DES1 for dihydroceramide substrate synthesized in the ER. All together, these data indicate that pathogenic SMS2 variants support bulk production of SM in the ER.

### Lipidome analysis of ER and PM isolated from cells expressing pathogenic SMS2 variants

We next asked whether pathogenic SMS2 variants that mediate bulk production of SM in the ER interfere with the ability of cells to generate a SM/cholesterol concentration gradient along the secretory pathway. To address this, we analyzed the lipid composition of ER and PM purified from wildtype or ΔSMS1/2 cells that express either SMS2 or the pathogenic variant SMS2^M64R^. For purification of the ER, cells were lysed and a post-nuclear supernatant was incubated with an antibody against calnexin (**Fig. 3a**). This was followed by incubation with secondary antibody-conjugated paramagnetic microbeads. For PM isolation, the surface of cells was treated with a non-membrane permeant biotinylation reagent before cell lysis (**Fig. 4a**). A post-nuclear supernatant was then directly incubated with streptavidin-conjugated paramagnetic microbeads. The microbeads were applied to columns packed with ferromagnetic spheres (μMACS columns) and the bound material was eluted after the columns were thoroughly washed. The purity of isolated ER and PM was assessed by immunoblot and lipidome analysis.

**Fig. 3.**
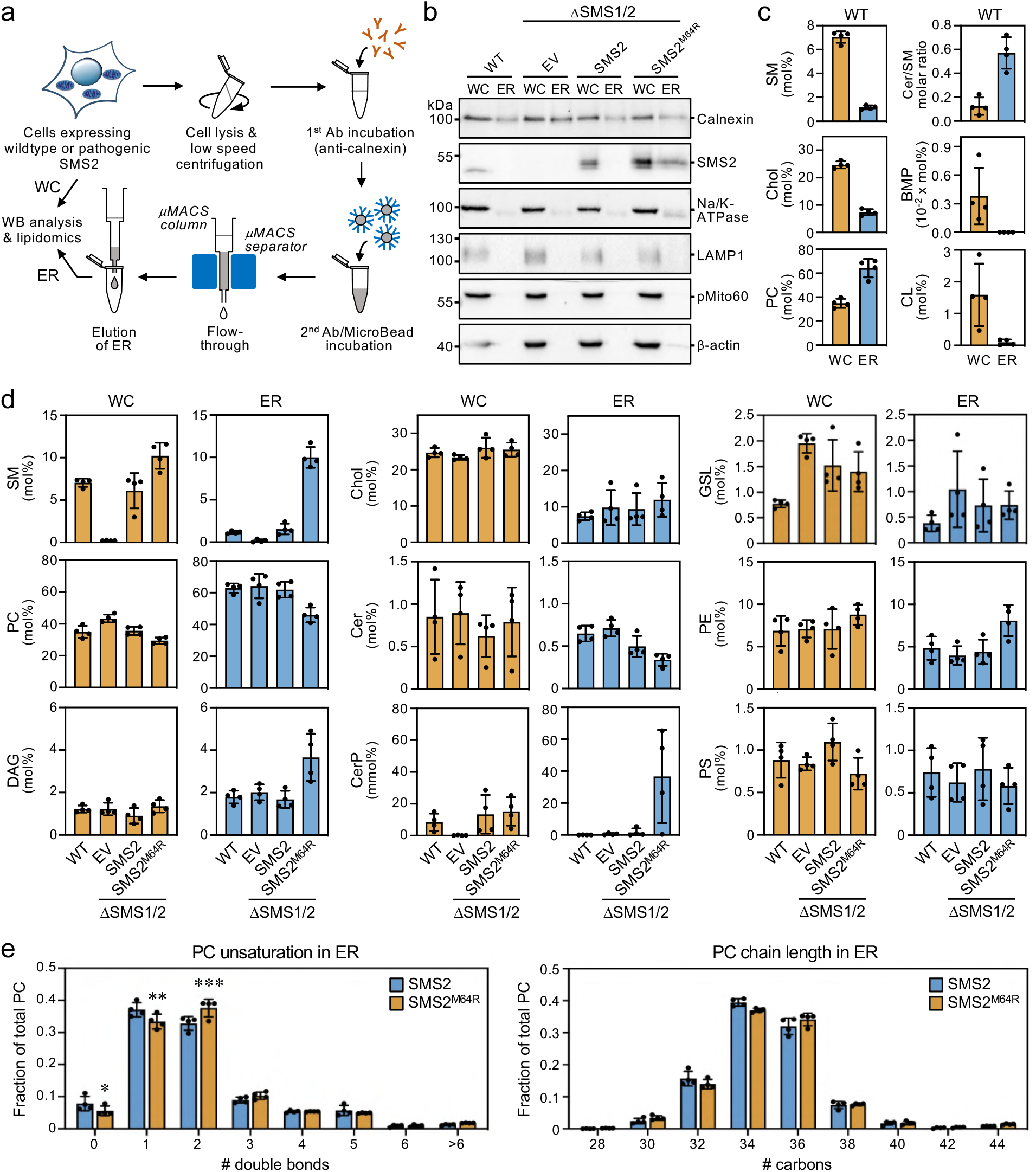
Cells expressing pathogenic variant SMS2^M64R^ accumulate SM in the ER. (**a**) Workflow for affinity purification of the ER from HeLa cells expressing wildtype or pathogenic SMS2 variants. (**b**) HeLa wildtype (WT) or ΔSMS1/2 cells transduced with empty vector (EV) or doxycycline-inducible SMS2 or SMS2^M64R^ were treated with doxycycline (1 μg/ml, 16 h), lysed and used to purify the ER as in (a). Whole cell lysates (WC) and purified ER were subjected to immunoblot analysis using antibodies against SMS2 and various organellar markers. (**c**) Lipid composition of whole cell lysates (WC) and ER purified from HeLa wildtype cells (WT) was determined by mass spectrometry-based shotgun lipidomics. Levels of the different lipid classes are expressed as mol% of total identified lipids. (**d**) Lipid composition of whole cell lysates (WC) and ER purified from cells as in (b) was determined as in (c). (**e**) Comparative analysis of PC unsaturation and chain length in ER purified from ΔSMS1/2 cells expressing SMS2 or SMS2^M64R^. The graphs show total numbers of double bonds (*left*) or carbon atoms (*right*) in the two acyl chains. All data are average ± SD, *n* = 4. \**p* < 0.05, \*\**p* < 0.01, \*\*\**p* < 0.001 by paired *t* test.

**Fig. 4.**
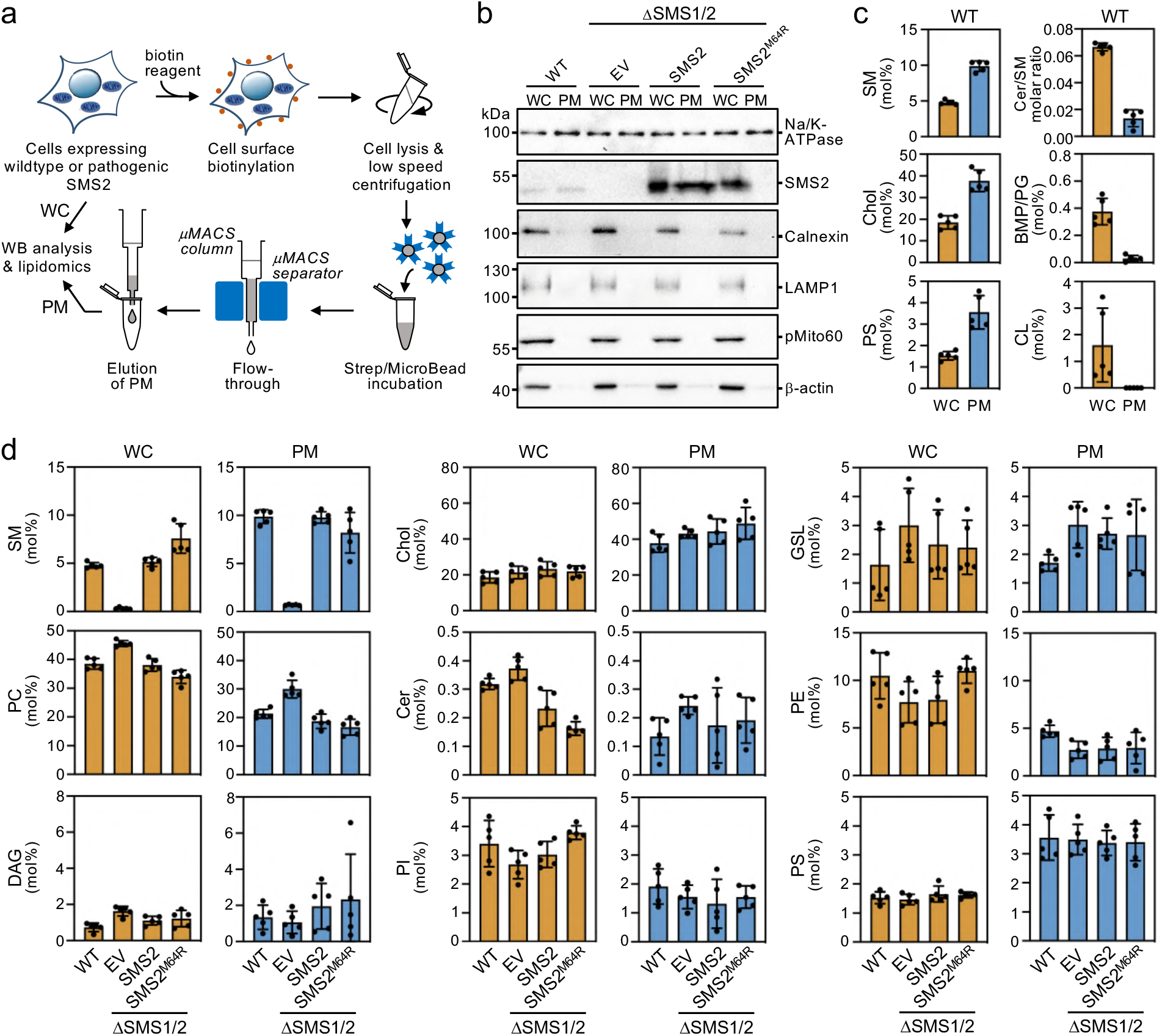
Lipid composition of the PM of cells expressing wildtype or pathogenic SMS2 variants. (**a**) Workflow for affinity purification of the PM from HeLa cells expressing wildtype or pathogenic SMS2 variants. (**b**) HeLa wildtype (WT) or ΔSMS1/2 cells transduced with empty vector (EV) or doxycycline-inducible SMS2 or SMS2^M64R^ were treated with doxycycline (1 μg/ml, 16 h), lysed and used to purify the PM as in (a). Whole cell lysates (WC) and purified PM were subjected to immunoblot analysis using antibodies against SMS2 and various organellar markers. (**c**) Lipid composition of whole cell lysates (WC) and PM purified from HeLa wildtype cells (WT) was determined by mass spectrometry-based shotgun lipidomics. Levels of the different lipid classes are expressed as mol% of total identified lipids. (**d**) Lipid composition of whole cell lysates (WC) and PM purified from cells as in (b) was determined as in (c). All data are average ± SD, *n* = 5.

As shown in **Fig. 3b**, ER eluates contained calnexin but were devoid of protein markers of the PM (Na/K-ATPase), lysosomes (LAMP1) or mitochondria (pMito60). As expected, ER eluates from cells expressing the pathogenic variant SMS2^M64R^ contained readily detectable levels of the protein. In contrast, no traces of SMS2 were found in ER eluates from cells expressing the wildtype protein. As there is no specific lipid marker for the ER, using a lipidomics approach to confirm that pull-down with anti-calnexin antibody indeed isolates the ER is not trivial. However, the ER is known to synthesize ceramides whereas SM is primarily produced in the *trans*-Golgi and accumulates in the PM. In line with the immunoblot data, ER eluates from wildtype cells displayed a 5-fold higher ceramide/SM ratio than total cell lysates. Moreover, ER eluates were largely devoid of lipids that are normally concentrated in mitochondria (cardiolipin, CL), PM (SM, cholesterol) and lysosomes (bis(monoacyl-glycerol)phosphate, here quantified together with the isobaric phosphatidylglycerol and reported as BMP/PG; **Fig. 3c**). Immunoblot analysis of the PM eluates revealed that they contain Na/K-ATPase but lack protein markers of the ER (calnexin), lysosomes (LAMP1) and mitochondria (pMito60; **Fig. 4b**). As expected, PM eluates from cells expressing wildtype SMS2 contained readily detectable amounts of the protein. On the other hand, PM eluates from ΔSMS1/2 cells expressing the pathogenic variant SMS2^M64R^ were devoid of this protein. Moreover, lipidome analysis of PM eluates revealed significantly elevated levels of lipids that are typically concentrated in the PM (i.e. SM, cholesterol, PS) and a 5-fold reduction in the ceramide/SM ratio relative to total cell lysates (**Fig. 4c**). Lipids primarily associated with lysosomes and mitochondria (BMP/PG, CL) were largely absent.

### Cells expressing pathogenic SMS2 variants accumulate SM in the ER

Using the pull-down approaches described above, we next determined the lipid composition of the ER and PM isolated from wildtype and ΔSMS1/2 cells expressing SMS2 or SMS2^M64R^. The ER from ΔSMS1/2 cells expressing SMS2 had a lipid composition similar to that of the ER from wildtype cells. In contrast, the ER from SMS2^M64R^-expressing cells contained 7-fold higher SM levels, i.e. ~10 mol% SM instead of ~1.5 mol% of all identified lipids (**Fig. 3d**). This increase in ER-bound SM was accompanied by a two-fold rise in DAG levels and a significant drop in the amount of PC and ceramide, consistent with the presence of a catalytically active SM synthase in the ER. Interestingly, expression of SMS2^M64R^ also led to a marked (1.8-fold) increase in ER-associated PE levels. In contrast, ER levels of cholesterol and other bulk lipids were largely unaffected. However, we noticed that expression of SMS2^M64R^ enhanced unsaturation of bulk phospholipid in the ER, as indicated by a significant rise in di-unsaturated PC at the expense of saturated and mono-unsaturated PC species (**Fig. 3e**). PC chain length, on the other hand, was not affected. Strikingly, SMS2^M64R^ expression also caused a sharp increase in ER-bound ceramide-1-phosphate (Cer1P). Moreover, cellular Cer1P levels were essentially abolished in SM synthase-deficient cells, indicating that production of Cer1P is tightly coupled to SM biosynthesis.

The PM from ΔSMS1/2 cells expressing SMS2 had a SM content similar to the PM from wildtype cells (~10 mol%). In comparison, the PM from ΔSMS1/2 cells expressing SMS2^M64R^ had a slightly reduced SM content (~8 mol%) even though the total SM content of these cells was considerably higher (**Fig. 4d**). PM-associated levels of cholesterol and other bulk lipids did not show any obvious differences among the various cell lines, except for an increase in PC and lack of SM in SMS-deficient cells. The PM from all four cell lines contained significantly elevated levels of saturated PC species in comparison to the ER. In addition, the PM from ΔSMS1/2 cells expressing SMS2^M64R^ contained 4-fold higher levels of dihydroSM (T. Sokoya, K. Maeda, and J. Holthuis, unpublished data), consistent with the ER residency of this enzyme. Collectively, these data indicate that pathogenic SMS2 variants disrupt the SM gradient along the secretory pathway and cause substantial changes in the lipid profile of the ER.

To confirm that cells expressing pathogenic SMS2 variants accumulate SM in the ER, we next used an engineered version of equinatoxin II (Eqt) as non-toxic SM reporter. To enable detection of SM inside the secretory pathway, the reporter was equipped with the *N*-terminal signal sequence of human growth hormone and tagged at its *C*-terminus with oxGFP, yielding EqtSM_SS_ (Deng et al., 2016). A luminal Eqt mutant defective in SM binding, EqtSol_SS_, served as control. When expressed in human osteosarcoma U2OS cells, both EqtSM_SS_ and EqtSol_SS_ showed a reticular distribution that overlapped extensively with the ER marker protein VAP-A (**Fig. 5a**). However, upon co-expression with SMS2^M64R^, EqtSM_SS_ but not EqtSol_SS_ displayed a distinct punctate distribution that coincided with the ER network. EqtSMSS-containing puncta were not observed upon co-expression with the enzyme-dead variant SMS2^M64R/D276A^, indicating that their formation strictly relies on SM production in the ER (**Fig. 5a**). To verify that the EqtSMSS-positive puncta mark ER-resident pools of SM, U2OS cells co-expressing EqtSM_SS_ and SMS2^M64R^ were subjected to hypotonic swelling as described before (King et al., 2020). After incubation for 5 min in hypotonic medium, the ER’s fine tubular network transformed into numerous micrometer-sized vesicles. In SMS2^M64R^-expressing cells, the membranes of these ER-derived vesicles were extensively labelled with EqtSM_SS_ (**Fig. 5b**). In contrast, in hypotonic cells co-expressing EqtSM_SS_ with SMS2^M64R/D276A^ or EqtSol_SS_ with SMS2^M64R^, the reporter was found exclusively in the lumen of ER-derived vesicles, indicating that Eqt staining of the ER membrane critically relies on catalytically active SMS2^M64R^ and a SM-binding competent reporter. In agreement with the ER lipidome analyses, these results demonstrate that cells expressing SMS2^M64R^ accumulate bulk amounts of SM in the ER. Moreover, our finding that hypotonic swelling of SMS2^M64R^-expressing cells transforms the ER-associated punctate distribution of EqtSM_SS_ to a more uniform labeling of the ER bilayer suggests that alterations in membrane curvature and/or lipid packing may affect the lateral organization of Eqt-SM assemblies.

**Fig. 5.**
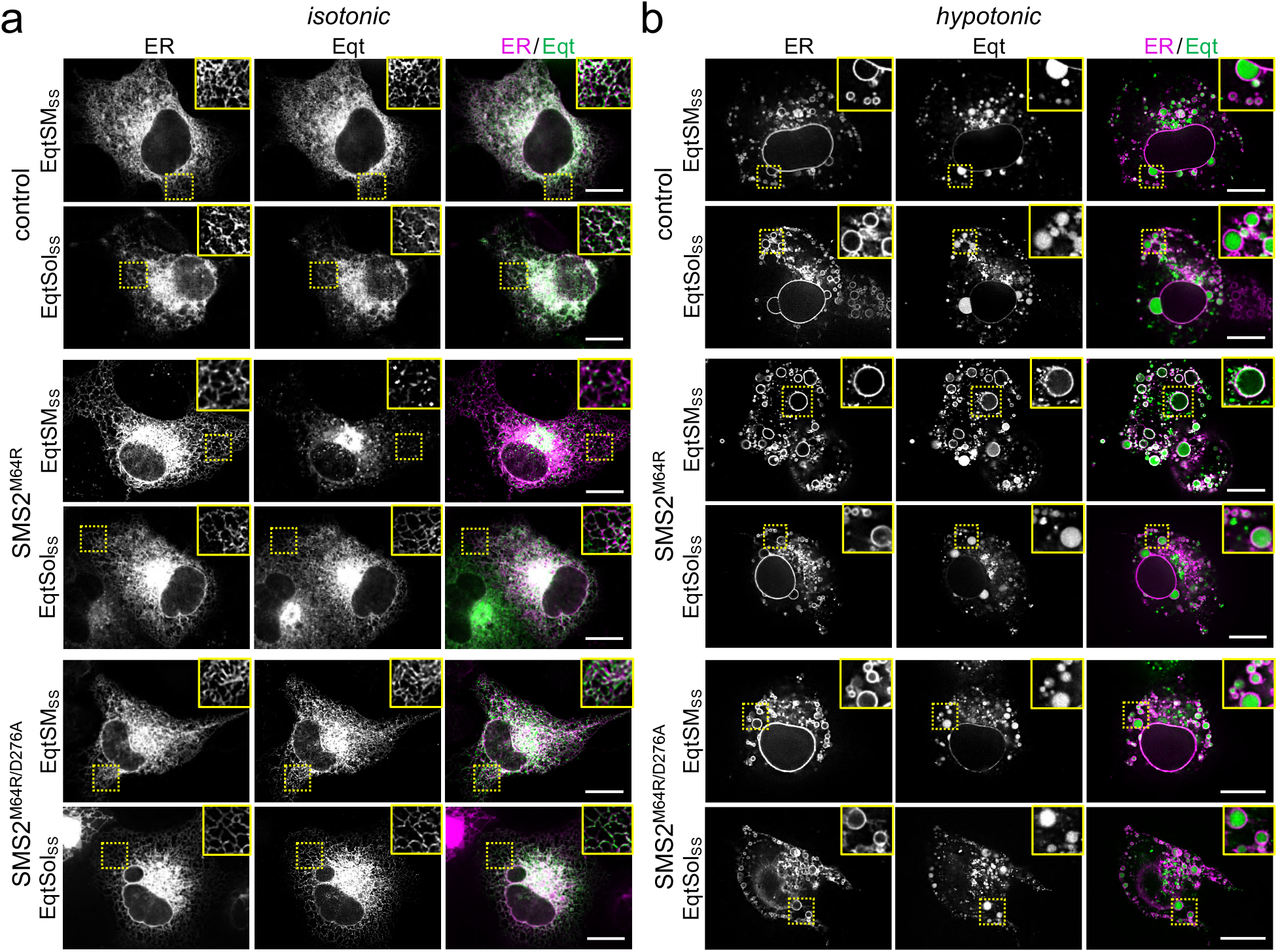
Luminal SM reporter EqtSMee enables visualization of an ER-resident SM pool in SMS2^M64R^-expressing cells. (**a**) Human osteosarcoma U2OS cells co-transfected with mCherry-tagged VAPA (ER, *magenta*) and empty vector (control), SMS2^M64R^ or SMS2^M64R/D276A^ and luminal GFP-tagged SM reporter EqtSM_SS_ or its SM binding-defective derivative, EqtSol_SS_ (Eqt, *green*), were incubated in isotonic medium (100% Optimem) for 5 min and imaged by spinning disc confocal microscopy. (**b**) Cells treated as in (a) were incubated in hypotonic medium (1% Optimem) for 5 min and then imaged by spinning disc confocal microscopy. Scale bar, 10 μm.

### Pathogenic SMS2 variants break transbilayer SM asymmetry

SM adopts an asymmetric distribution across the bilayers of late secretory and endolysosomal organelles, with the bulk of SM residing in the luminal/exoplasmic leaflet. However, using GFP-tagged EqtSM as cytosolic SM reporter (EqtSM_cyto_), we found that perturbation of lysosome or PM integrity by pore-forming chemicals or toxins disrupts transbilayer SM asymmetry by triggering a rapid transbilayer movement of SM catalyzed by Ca^2+^-activated scramblases (Niekamp et al., 2022). To perform its central task in membrane biogenesis, the ER harbors constitutively active scramblases that enable a rapid equilibration of newly synthesized phospholipids across its bilayer (Pomorski and Menon, 2016). We therefore asked whether SM produced by pathogenic SMS2 variants in the ER lumen has access to the cytosolic leaflet. As expected, EqtSM_cyto_ in wildtype or ΔSMS1/2 cells expressing SMS2 displayed a cytosolic distribution. In contrast, expression of pathogenic variant SMS2^I62S^ or SMS2^M64R^ in each case caused EqtSM_cyto_ to accumulate in numerous puncta that were dispersed throughout the cytosol (**Fig. 6a**; **Fig. S3a**). Importantly, formation of Eqt-positive puncta required expression of a catalytically active pathogenic variant and was not observed when using SM binding-defective cytosolic reporter EqtSol_cyto_ (**Fig. S3b**). These results indicate that pathogenic SMS2 variants disrupt transbilayer SM asymmetry, presumably because ER-resident scramblases enable SM produced by these variants to readily equilibrate across the ER bilayer.

**Fig. 6.**
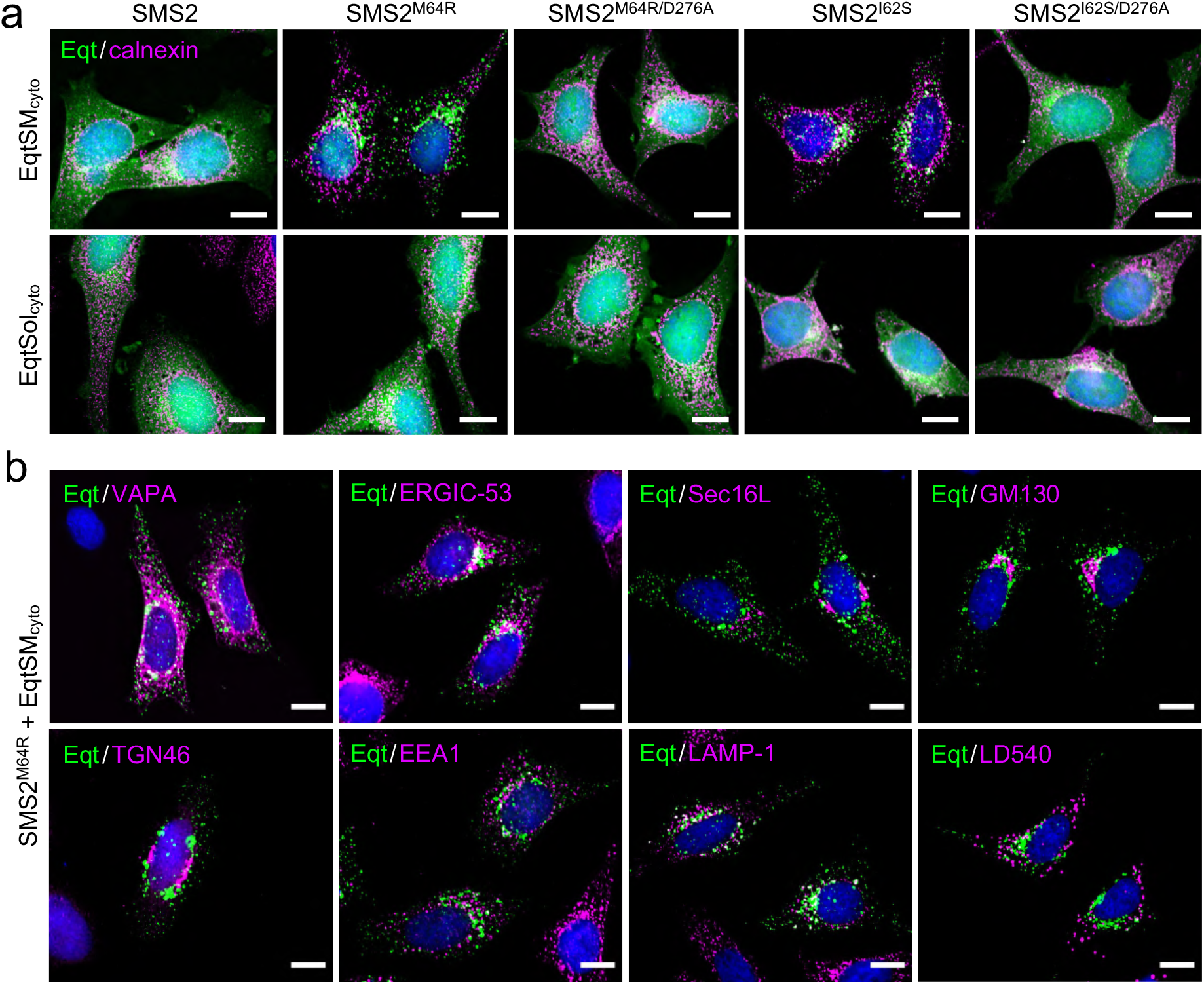
Pathogenic SMS2 variants disrupt transbilayer SM asymmetry. (**a**) HeLa ΔSMS1/2 cells transduced with doxycycline-inducible SMS2, SMS2^M64R^, SMS2^I62S^ or their enzyme-dead isoforms (D276A) were transfected with cytosolic GFP-tagged SM reporter EqtSMcyto or its SM binding-defective derivative, EqtSolcyto (Eqt, *green*). After treatment with doxycycline (1 μg/ml, 16 h), cells were fixed, immunostained with a-calnexin antibodies (*magenta*), counterstained with DAPI (*blue*) and imaged by DeltaVision microscopy. (**b**) HeLa ΔSMS1/2 cells transduced with doxycycline-inducible SMS2^M64R^ were transfected with EqtSMcyto (*green*) and treated with doxycycline as in (a). Next, cells were fixed, immunostained with antibodies against various organellar markers (*magenta*), counterstained with DAPI (*blue*) and imaged by DeltaVision microscopy. The ER was marked by co-transfection with mCherry-tagged VAPA while lipid droplets were labeled using the lipophilic dye LD540. Scale bar, 10 μm.

Remarkably, the Eqt-positive puncta formed in SMS2^I62S^ or SMS2^M64R^-expressing cells were largely segregated from a wide array or organellar markers, including VAPA (ER), Sec16L (ER exit sites), ERGIC-53 (*cis*-Golgi), GM130 (*medial*-Golgi), TGN64 (*trans*-Golgi), EEA1 (early endosomes), LAMP1 (lysosomes) and LD540 (lipid droplets; **Fig. 6b**). Moreover, when cells expressing SMS2^I62S^ or SMS2^M64R^ were subjected to hypotonic swelling, Eqt-positive puncta remained largely segregated from ER-derived microvesicles (T. Sokoya and J. Holthuis, unpublished data). Conceivably, formation of Eqt-SM assemblies on the cytosolic surface of the ER may drive a process whereby SM-rich membrane domains are pinched off from the organelle. However, our efforts to capture such vesicles by correlative light-electron microscopy were unsuccessful. Therefore, the precise nature of the Eqt-positive puncta observed in cells expressing pathogenic variants remains to be established.

The foregoing implies that in cells expressing pathogenic SMS2 variants, only part of the SM arriving at the PM would reside in the exoplasmic leaflet and that a portion may be mislocalized to the cytosolic leaflet. To challenge this idea, we stained the surface of intact wildtype or ΔSMS1/2 cells expressing SMS2 or SMS2^M64R^ with recombinant EqtSM. Cell surface labeling was visualized by fluorescence microscopy and quantitatively assessed by flow cytometry. As shown in **Fig. 7a**, wildtype cells could be readily stained with the SM reporter whereas ΔSMS1/2 cells were devoid of EqtSM staining. Expression of SMS2, but not enzyme-dead SMS2^D276A^, restored EqtSM staining of ΔSMS1/2 cells to a level approaching that of wildtype cells. In contrast, expression of SMS2^M64R^ restored cell surface staining to only a minor degree (**Fig. 7a, b**) even though the PM-associated SM pool of these cells was close to that of wildtype or SMS2-expressing ΔSMS1/2 cells (**Fig. 4d**). Collectively, these data suggest that pathogenic SMS2 variants undermine the ability of cells to establish transbilayer SM asymmetry.

**Fig. 7.**
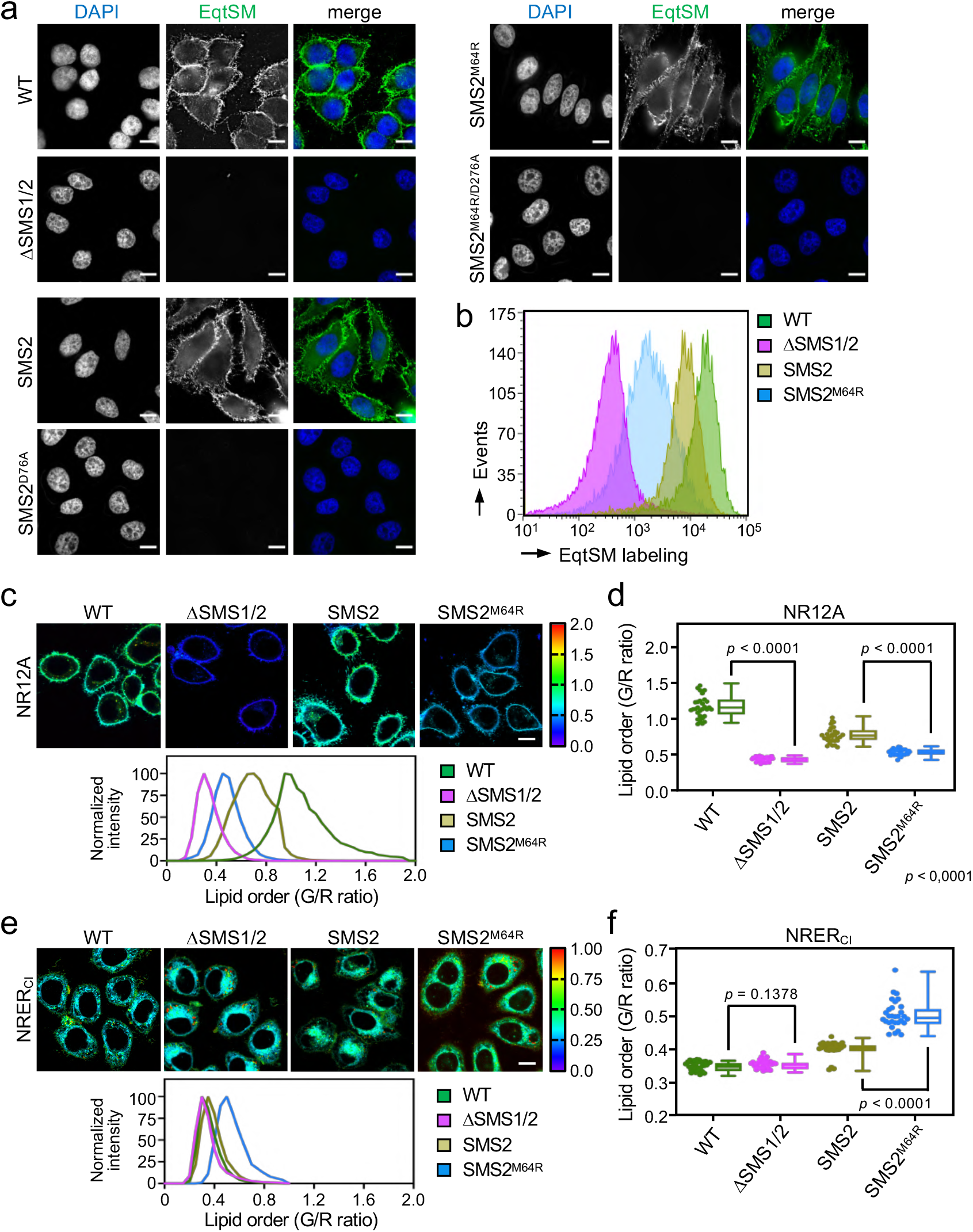
SMS2^M64R^-expressing cells fail to concentrate SM on their surface and exhibit imbalances in lipid order. (**a**) HeLa wildtype (WT) or ΔSMS1/2 cells transduced with doxycycline-inducible SMS2, SMS2^M64R^ or their enzyme-dead isoforms (D276A) were treated with doxycycline (1 μg/ml, 16 h), incubated with FLAG-tagged EqtSM, fixed, co-stained with a-FLAG antibody (*green*) and DAPI (*blue*), and imaged by DeltaVision microscopy. (**b**) Cells treated as in (a) were analyzed by flow cytometry to quantitatively assess EqtSM labeling of their surface. (**c**) Cells treated as in (a) were stained with 0.2 μM NR12A for 10 min and analyzed by ratiometric fluorescence microscopy to probe the lipid order in the outer PM leaflet. Warmer colors reflect a higher lipid order. (**d**) Quantitative assessment of changes in lipid order in the outer PM leaflet of cells treated as in (c). *n* = 30 cells per condition over two independent experiments. (**e**) Cells treated as in (a) were stained with 0.2 μM NRER_CI_ for 10 min and analyzed by ratiometric fluorescence microscopy to probe lipid order in the ER. Warmer colors reflect a higher lipid order. (**f**) Quantitative assessment of changes in lipid order in the ER of cells treated as in (e). *n* = 30 cells per condition over two independent experiments. All *p* values calculated by unpaired *t*-test. Scale bar, 10 μm.

### Pathogenic SMS2 variants affect membrane lipid order along the secretory pathway

Owing to its saturated nature and affinity for sterols, SM contributes significantly to lipid order in cellular membranes. As pathogenic SMS2 variants enhance SM levels in the ER and undermine the ability of cells to concentrate SM on their surface, we next measured the lipid order at these locations in cells expressing SMS2 or SMS2^M64R^ using two Nile Red (NR)-based solvatochromic probes, NR12A and NRER_CI_ (Danylchuk et al., 2019, 2021). The emission spectra of these probes are blue-shifted in tightly packed lipid bilayers, which is due to a reduced polarity in the nano-environment of the NR fluorophore. The relative extent of these spectral shifts can be quantified in cellular membranes using ratiometric imaging. In NR12A, the presence of a charged membrane anchor group that blocks passive flip-flop across membrane bilayers makes this probe ideally suited to selectively quantify lipid order in the outer PM leaflet of live cells when added to the extracellular medium (Danylchuck et al., 2019). On the other hand, a propyl chloride group in NRER_CI_ targets this probe to the ER (Danylchuck et al., 2021). As expected, the lipid order reported by NR12A in the outer PM leaflet of SM-deficient ΔSMS1/2 cells was drastically reduced in comparison to that of wildtype cells (**Fig. 7c, d**). Expression of SMS2 partially restored lipid order. In contrast, expression of SMS2^M64R^ failed to restore lipid order to any appreciable degree. Conversely, the lipid order reported by NRER_CI_ in the ER of SMS2^M64R^-expressing cells was significantly enhanced in comparison to that in the ER of SMS2-expressing cells (**Fig. 7e, f**). Hence, the perturbation of subcellular SM distributions caused by pathogenic SMS2 variants is accompanied by major imbalances in lipid order along the secretory pathway.

### Pathogenic SMS2 variants perturb subcellular cholesterol pools

As preferred cholesterol interaction partner, SM directly participates in the subcellular organization of cholesterol (Slotte, 2013; Das et al., 2014). Therefore, it was surprising that the marked accumulation of SM in the ER of SMS^M64R^-expressing cells had no obvious impact on ER-bound cholesterol levels (**Fig. 3d**) or the cellular pool of cholesteryl esters (T. Sokoya, K. Maeda, and J. Holthuis, unpublished data). To further examine whether pathogenic SMS2 variants influence cholesterol organization in cells, we used a mCherry-tagged D4H sterol reporter derived from the perfringolysin O *θ*-toxin of *Clostridium perfringens*. This reporter recognizes cholesterol when present at >20 mol% in membranes and mainly decorates the inner leaflet of the PM at steady state when expressed as cytosolic protein. Part of the reason for the high detection threshold is that the reporter detects free cholesterol in the membrane but not cholesterol in complex with SM (Das et al., 2014). Moreover, a previous study showed that plasmalemmal PS is essential for retaining D4H-accessible cholesterol in the inner PM leaflet (Maekawa and Fairn, 2015). Accordingly, we found that cytosolic D4H-mCherry primarily stained the PM in wildtype HeLa cells. In contrast, the probe displayed a more diffuse cytosolic distribution in ΔSMS1/2 cells (**Fig. S4a**). Expression of SMS2 in ΔSMS1/2 cells restored PM staining, indicating that PM-associated SM is critical for controlling the reporter-accessible cholesterol pool in the PM. Strikingly, in ΔSMS1/2 cells expressing pathogenic variant SMS2^M64R^ or SMS2^I62S^, D4H-mCherry did not label the PM but primarily accumulated in intracellular vesicles that were largely segregated from EqtSM_cyto_-positive puncta (**Fig. 8a**). These vesicles co-localized extensively with dextran-positive endolysosomal compartments (**Figs. 8b** and **S4b**). While it remains to be established how this shift in D4H-mCherry distribution is accomplished, it is conceivable that the presence of SM in the inner PM leaflet of cells expressing pathogenic SMS2 variants renders a coexisting cholesterol pool inaccessible to the reporter.

**Fig. 8.**
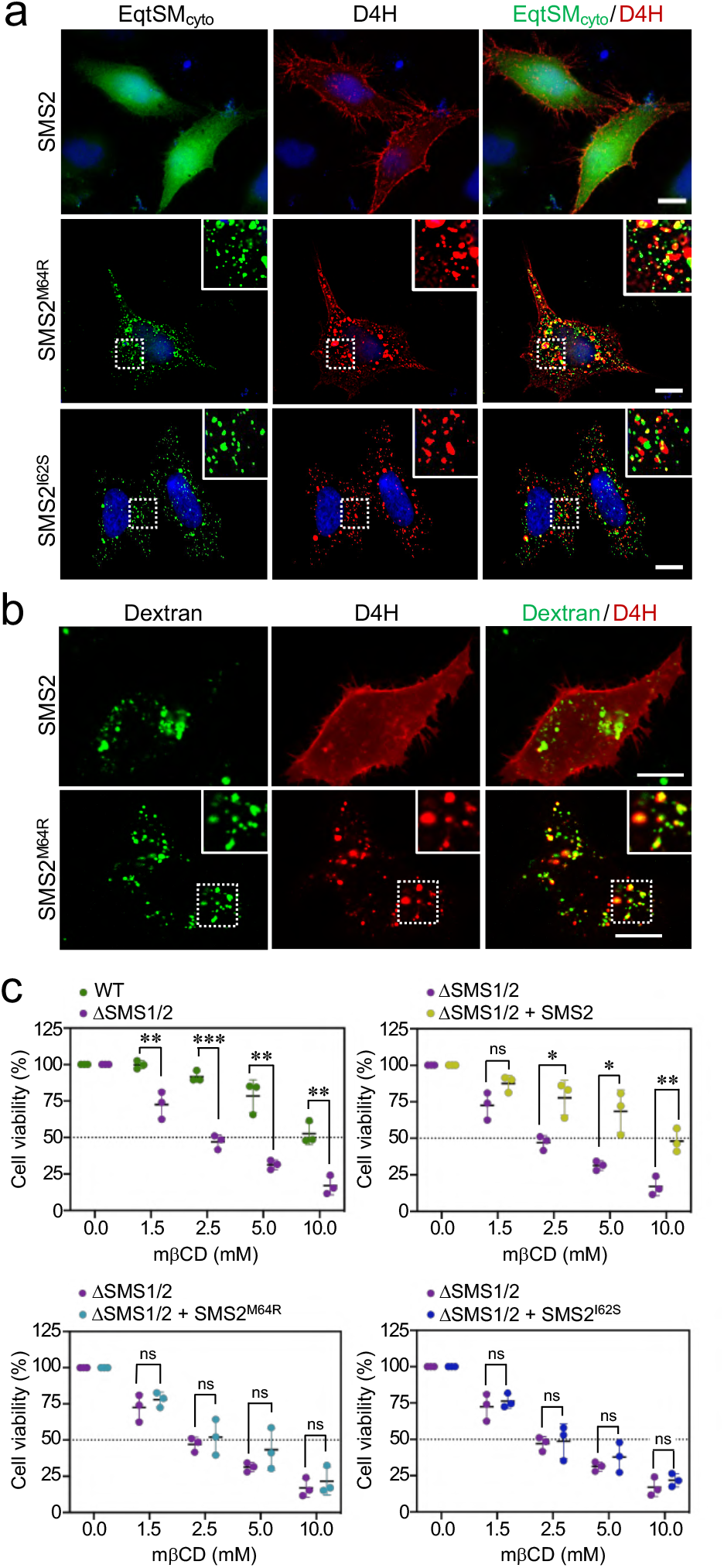
Pathogenic SMS2 variants perturb subcellular cholesterol pools. (**a**) HeLa ΔSMS1/2 cells transduced with doxycycline-inducible SMS2, SMS2^M64R^ or SMS2^I62S^ were cotransfected with GFP-tagged EqtSMcyto (*green*) and mCherry-tagged cytosolic sterol reporter D4H (*red*). Next, cells were treated with 1 μg/ml doxycycline for 16 h, fixed, counterstained with DAPI (*blue*) and visualized by DeltaVision microscopy. (**b**) HeLa ΔSMS1/2 cells stably transduced with FLAG-tagged SMS2 or SMS2^M64R^ were transfected with mCherry-tagged D4H (*red*), labeled with fluorescein-conjugated dextran (*green*) in the presence of 1 μg/ml doxycycline for 16 h and then imaged by spinning disc confocal microscopy. (**c**) HeLa wildtype (WT) or ΔSMS1/2 cells stably transduced with SMS2, SMS2^M64R^ or SMS2^I62S^ were treated with 1 μg/ml doxycycline for 16 h. Next, cells were exposed to the indicated concentration of methyl β-cyclodextrin (mβCD) for 1 h and cell viability was assessed using Prestoblue reagent. Data shown are averages of 4 technical replicates from *n* = 3 biological replicates. \**p* < 0.05, \*\**p* < 0.01, \*\*\**p* < 0.001 by paired *t* test. Scale bar, 10 μm.

Since cholesterol has a stronger affinity for SM than for PS or other phospholipid classes, altering the SM concentration affects the behavior of cholesterol in artificial and biological membranes (Slotte, 2013). For instance, when exposed to the cholesterol-absorbing agent methyl-β-cyclodextrin (mβCD), SM-depleted cells readily lose PM-associated cholesterol and consequently their viability more rapidly than wildtype cells (Fukasawa et al., 2000; Hanada et al., 2003). As complementary approach to determine the impact of pathogenic SMS2 variants on cholesterol organization in the PM, we next probed ΔSMS1/2 cells expressing wildtype or pathogenic SMS2 variants for their sensitivity toward mβCD. As expected, ΔSMS1/2 cells displayed a substantially reduced tolerance for mβCD in comparison to wildtype cells (**Fig. 8c**). Expression of SMS2 restored mβCD tolerance of ΔSMS1/2 cells to that of wildtype cells. In contrast, expression of SMS^M64R^ or SMS2^I62S^ in each case failed to render ΔSMS1/2 cells resistant toward mβCD. These results provide additional support for the notion that pathogenic SMS2 variants significantly affect cholesterol organization in the PM.

### Patient-derived fibroblasts display imbalances in SM distribution and lipid packing

We next asked whether the aberrant SM and cholesterol distributions observed upon heterologous expression of pathogenic SMS2 variants also occur in cells of patients with OP-CDL. To address this, skin fibroblasts derived from patients with the missense variant p.I62S or p.M64R and healthy controls were co-transfected with the luminal SM reporter EqtSM_SS_ and mCherry-tagged VAPA as ER marker. Next, the fibroblasts were subjected to hypotonic swelling and imaged by fluorescence microscopy. Strikingly, in patient fibroblasts the membranes of ER-derived vesicles were extensively labelled with EqtSM_SS_ (**Fig. 9a**). In contrast, in fibroblasts of healthy controls, EqtSM_SS_ was found exclusively in the lumen of ER-derived vesicles. This indicates that the ER in patient fibroblasts contains substantially elevated SM levels. In addition, we found that the cytosolic SM reporter EqtSM_cyto_ accumulated in numerous puncta when expressed in patient fibroblasts while its expression in fibroblasts of healthy controls resulted in a diffuse cytosolic distribution (**Fig. 9b**). Formation of Eqt-positive puncta in patient fibroblasts did not occur when using the SM binding-defective reporter, EqtSol_cyto_. Thus, besides accumulating SM in the ER, patient fibroblasts display a breakdown of transbilayer SM asymmetry. Interestingly, these aberrant SM distributions were accompanied by significant alterations in lipid order on the cell surface (**Fig. 9c**) and in the ER (**Fig. 9d**). Moreover, while the cytosolic cholesterol reporter D4H-mCherry primarily stained the PM of fibroblasts of healthy controls, in patient fibroblasts a substantial portion of the reporter was shifted to intracellular vesicles (**Fig. S5**). This indicates that pathogenic SMS2 variants p.I62S and p.M64R, which underly a spectrum of severe skeletal conditions, affect the subcellular organization of SM and cholesterol to an extend large enough to impact on the lipid order along membranes of the secretory pathway.

**Fig. 9.**
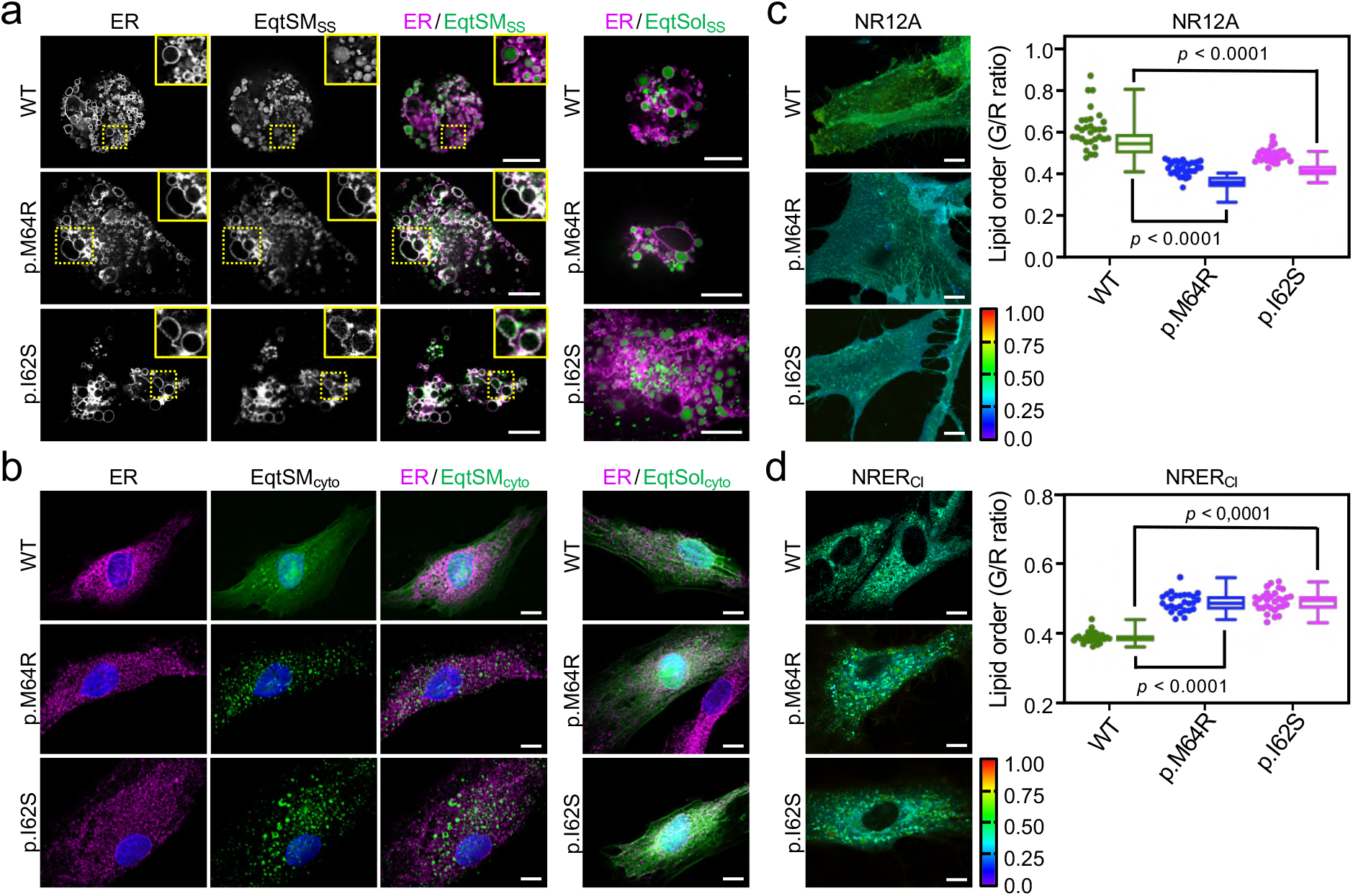
Patient-derived fibroblasts display perturbations in SM distribution and lipid order. (**a**) Control (WT) or patient-derived human skin fibroblasts carrying heterozygous missense variants c.185T>G (p.I62S) or c.191T>G (p.M64R) in *SGMS2* were co-transfected with mCherry-tagged VAPA (ER, *magenta*) and GFP-tagged EqtSM_SS_ (*green*). Co-transfections with GFP-tagged EqtSol_SS_ served as control. After 16 h, cells were incubated in hypotonic medium (1% Optimem) for 5 min and imaged by spinning disc confocal microscopy. (**b**) Fibroblasts as in (a) were transfected with GFP-tagged EqtSMcyto. After 16 h, cells were fixed, immunostained with a-calnexin antibodies (ER, *magenta*), counterstained with DAPI (*blue*) and imaged by DeltaVision microscopy. (**c**) Fibriblasts as in (a) were stained with 0.2 μM NR12A for 10 min and analyzed by ratiometric fluorescence microscopy to quantitatively access lipid order in the outer PM leaflet. Warmer colors reflect a higher lipid order. *n* = 30 cells per condition analyzed over two independent experiments. (**d**) Fibroblasts as in (a) were stained with 0.2 μM NRER_CI_ for 10 min and analyzed by ratiometric fluorescence microscopy to quantitatively access lipid order in the ER. Warmer colors reflect a higher lipid order. *n*= 27 cells per condition analyzed over two independent experiments. All *p* values calculated by unpaired *t*-test. Scale bar, 10 μm.

## DISCUSSION

SM in mammalian cells is specifically enriched in the exoplasmic leaflets of the PM, the *trans*-Golgi and endolysosomal organelles. While maintenance of its nonrandom subcellular distribution is thought to be relevant for a variety of physiological processes, experimental proof for this concept is scarce. Here we show that inborn pathogenic SMS2 variants p.M64R and p.I62S identified in patients with a severe form of OP-CDL cause profound perturbations in the subcellular organization of SM and cholesterol. We find that both variants retain full enzymatic activity but are unable to leave the ER owing to a defective autonomous ER export signal in their *N*-terminal cytosolic tails. Consequently, bulk SM production is mistargeted to the ER, the site for *de novo* synthesis of the SM precursor ceramide. Cells expressing pathogenic SMS2 variants accumulate PM-like SM levels in the ER and display a disrupted transbilayer SM asymmetry, presumably owing to a constitutive SM scrambling across the ER bilayer. These aberrant SM distributions also occur in OP-CDL patient fibroblasts and are accompanied by significant imbalances in cholesterol organization, glycerophospholipid profiles and membrane lipid order in the secretory pathway. Based on these findings, we postulate that pathogenic SMS2 variants undermine the capacity of osteogenic cells to uphold nonrandom lipid distributions that are critical for their bone forming activity.

As SM is the preferred interaction partner of cholesterol (Slotte, 2013), its bulk production in the *trans*-Golgi would promote formation of a cholesterol gradient along the secretory pathway. However, our lipidomics data indicate that cells harboring a pathogenic SMS2 variant retain the ability to concentrate cholesterol in the PM and keep its ER levels low in spite of a dissipated SM gradient. This implies that cells are equipped with an effective mechanism to prevent a toxic rise of cholesterol in ER bilayers with an abnormally high SM content. One mechanism for removing excess cholesterol from the ER involves its esterification and storage in lipid droplets. However, pathogenic SMS2 variants had no impact on the cellular pool of cholesteryl esters. An alternative mechanism involves the oxysterol binding protein OSBP, which mediates net transfer of cholesterol from the ER to the *trans*-Golgi. OSBP-catalyzed transport of cholesterol against its concentration gradient is energized by counter transport of phosphatidylinositol-4-phosphate (PI4P), a lipid continuously produced in the *trans*-Golgi and turned over in the ER (Mesmin et al., 2013). As critical determinant of intracellular cholesterol flows, the cholesterol/PI4P exchange activity of OSBP also influences membrane lipid order in the secretory pathway (Mesmin et al., 2017). Therefore, future studies addressing whether pathogenic SMS2 variants cause an upregulation of the PI4P-consuming OSBP cycle to counteract thermodynamic trapping of cholesterol by an expanding SM pool in the ER may proof fruitful.

The PI4P-dependent countertransport mechanism found to energize net transfer of cholesterol also helps drive PS export from the ER and creation of a PS gradient along the secretory pathway (Chung et al., 2015; Moser von Filseck et al., 2015). Our current findings indicate that such mechanism does not exist for SM and that the ER is ill equipped to effectively remove bulk amounts of SM produced there. In the absence of any lipid transfer protein dedicated to mediate anterograde SM transport, the only mechanism available for SM to leave the ER is by vesicular transport. Previous work revealed that SM and cholesterol are depleted from COPI-coated vesicles compared with their donor Golgi cisternae (Brügger et al., 2000). This finding supports the view that formation of the SM gradient along the secretory pathway relies on a mechanism that prevents *trans*-Golgi-derived SM from gaining access to retrograde-moving COPI vesicles. Experiments with giant unilamellar vesicles containing ternary mixtures of PC, SM and cholesterol provide evidence for curvature-based lipid sorting by demonstrating that membrane tubes pulled from the giant vesicles are efficiently depleted of SM and cholesterol relative to vesicle membranes with negligible curvature (Roux et al., 2005). The same principle may hamper an efficient COPII-mediated export of SM from the ER, thus contributing to dissipation of the SM gradient in cells harboring pathogenic SMS2 variants.

Membrane biogenesis in the ER requires cross-bilayer movement of phospholipids, which is mediated by ER-resident scramblases (Pomorski and Menon, 2016; Ghanbarpour et al., 2021; Huang et al., 2021). These scramblases display low specificity, with phospholipids and sphingolipids being translocated with similar kinetics (Buton et al., 2002; Chalat et al., 2012). Consequently, SM produced by pathogenic SMS2 variants in the luminal leaflet of the ER should readily equilibrate with the cytosolic leaflet. Indeed, we found that both p.M64R and p.I62S variants triggered mobilization of a cytosolic SM reporter. Moreover, cells expressing the p.M64R variant had similar levels of SM in the PM as controls but showed a significantly reduced SM reporter staining of their surface, signifying a disrupted transbilayer SM asymmetry. Consistent with a reduced SM pool in the exoplasmic leaflet, these cells also displayed a lower lipid packing on their surface and lost their viability more rapidly when exposed to a cholesterol-absorbing agent than controls. Consistent with an elevated SM pool in the cytosolic leaflet, we found that pathogenic SMS2 variants constrained accessibility of PM-associated cholesterol for a cytosolic sterol reporter that recognizes free cholesterol but not cholesterol sequestered by SM (Das et al., 2014). From this we infer that pathogenic SMS2 variants in fact abolish two types of SM gradients: one running along the secretory pathway and the other one across the bilayers of secretory organelles. Disruption of the latter affects the equilibrium between active and SM-sequestered cholesterol pools on both sides of the PM.

Our study also yields insights into how cells cope with a major assault on the lipid composition and membrane properties of the ER. The dramatic rise in ER-associated SM levels caused by pathogenic SMS2 variants was accompanied by a marked increase in PC desaturation and a nearly two-fold expansion of the ethanolamine-containing phospholipid pool. While it remains to be established how these alterations are implemented, it is conceivable that they serve to buffer the physical properties of the ER bilayer from SM-induced perturbations. Owing to the low degree of unsaturation in its carbohydrate chains, SM forms a taller, narrower cylinder than PC, which increases its packing density and affinity for cholesterol (Slotte, 2013). These features are ideally suited to support the barrier function of the PM but undermine the biogenic activities of the ER, which require a more loosely packed lipid bilayer (Bigay and Antonny, 2012; Nilsson et al., 2001). Thus, an enhanced desaturation and rise in cone-shaped, ethanolamine-containing phospholipids may be part of an adaptive cellular response to cancel out a SM-mediated rigidification of the ER bilayer and preserve the organelle’s central role in membrane biogenesis and secretion. Strikingly, pathogenic SMS2 variants also caused a sharp rise in ER-associated Cer1P levels. Cer1P is produced by ceramide kinase CERK and functions as a key signaling lipid in the regulation of cell growth, survival and inflammation (Presa et al., 2020). In addition to stimulating production of arachidonic acid and pro-inflammatory cytokines through direct activation of a specific cytosolic phospholipase (Lamour et al., 2009), Cer1P promotes cell survival at least in part by blocking enzymes involved in ceramide production (Granado et al., 2009). Our present findings indicate that Cer1P production is tightly coupled to SM biosynthesis. The prospect that CERK-mediated Cer1P formation serves a role in the mechanism by which cells sense and respond to imbalances in the lipid composition of their secretory organelles merits further consideration.

How does a disrupted subcellular organization of SM and cholesterol caused by pathogenic SMS2 variants lead to osteoporosis and skeletal dysplasia? Addressing this question obviously requires experimental models beyond the engineered cell lines and patient-derived fibroblasts used in this study. A quantitative analysis of SMS2 transcript levels in a murine tissue panel revealed the highest expression in cortical bone and vertebrae (Pekkinen et al., 2019). This implies that the impact of pathogenic SMS2 variants on the lipid composition of secretory organelles will be most severe in bone cells of the affected individuals. Bone formation involves deposition of collagen fibrils into a matrix and its subsequent mineralization. Interestingly, pathogenic variants of core components of COPII-coated vesicles have been reported to cause craniofacial and skeletal defects by selectively disrupting procollagen export from the ER (Boyadjiev et al., 2006; Garbes et al., 2015). Moreover, loss of TANGO1, an ER-resident transmembrane protein required for packaging the bulky procollagen fibers into COPII vesicles, results in neonatal lethality due to insufficient bone mineralization (Guillemyn et al., 2021). It is conceivable that ER export of procollagen is particularly susceptible to the bilayer rigidifying effect of bulk SM production by pathogenic SMS2 variants.

However, an alternative scenario is that a loss of transbilayer SM asymmetry at the PM of osteogenic cells in OP-CDL patients negatively affects bone mineralization. This process involves matrix vesicles that bud off from the apical membrane of osteoblasts and deposit their Ca^2+^ and phosphate-rich content where matrix mineralization is propagated (Murshed, 2018). Bone mineralization also critically relies on neutral SMase-2 (SMPD3), a membrane-bound enzyme that cleaves SM in the cytosolic leaflet of the PM to generate ceramide and phosphocholine (Aubin et al., 2005). How nSMase-2 gains access to SM, which is normally concentrated in the exoplasmic leaflet, is unclear. We recently showed that minor lesions in PM integrity initiates a rapid SM scrambling mediated by the Ca^2+^-activated scramblase TMEM16F (Niekamp et al., 2022). Intriguingly, loss of TMEM16F leads to decreased mineral deposition in skeletal tissues (Ehlen et al., 2013), suggesting that this process may require a TMEM16F-mediated supply of exoplasmic SM to nSMase-2 for SM hydrolysis in the cytosolic leaflet. This arrangement may serve to ensure a continuous supply of phosphocholine as a source of phosphate required for normal bone mineralization (Pekkinen et al., 2019). By disrupting SM asymmetry, pathogenic SMS2 variants may cause premature depletion of the lipid-based phosphate store, thus interfering with normal bone mineralization. Moreover, ceramides formed during SM hydrolysis by nSMase-2 in the cytosolic leaflet of the PM may also play a critical role. Owing to their cone-shaped structure, ceramides released by SM turnover readily self-assemble into microdomains that possess a negative spontaneous curvature (Alonso and Goñi, 2018). By causing a local condensation of the cytosolic leaflet, nSMase-mediated conversion of SM to ceramide could promote an inverse budding of the bilayer away from the cytosol. We envision that this process may stimulate the biogenesis of matrix vesicles required for normal bone mineralization. According to this model, pathogenic SMS2 variants may interfere with matrix vesicle formation because dissipation of transbilayer SM asymmetry across the PM would deplete the fuel that drives this process.

In sum, the present study indicates that bone critical SMS2 variants p.M64R and p.I62S exert their pathogenic effects by redirecting bulk SM production to the ER, thereby causing significant deviations in organellar lipid compositions and membrane properties along the secretory pathway. Besides highlighting how cells respond to a major assault on the lipid code of the early secretory pathway, our findings provide important insights into the pathogenic mechanism underlying OP-CDL.

## METHODS

### Chemical reagents

Chemical reagents used were: doxycycline (Sigma Aldrich, D891), puromycin (Sigma Aldrich, P8833), polybrene (Sigma-Aldrich, TR-1003), methyl-β-cyclodextrin (Sigma Aldrich, C4555), G418 (Sigma-Aldrich, G8168), and 3-azido-7-hydroxycoumarin (Jena Bioscience, CLK-FA047). LD540 dye was a kind gift from Christoph Thiele (University of Bonn, Germany) and described in (Spandl et al., 2009). NR12A and NR-ERcl were synthesized as described in (Danylchuk et al., 2021).

### Antibodies

Antibodies used were: rabbit polyclonal anti-calnexin (Santa Cruz, sc-11397; IB 1:1,000), goat polyclonal anti-calnexin (Santa Cruz, sc-6495; IF 1:200, IB 1:1,000), rabbit polyclonal anti-β-calnexin (Abcam, ab10286; IP 1:1,000), mouse monoclonal anti-FLAG-tag (Abcam, ab205606; IB 1:1,000; IF 1:400), mouse monoclonal anti-SMS2 (Santa Cruz, sc-293384; IB 1:1,000), mouse monoclonal anti-β-actin (Sigma, A1978; IB 1:50,000), mouse monoclonal anti-mitochondrial surface p60 (Millipore, MAB1273; IB 1:1,000), mouse monoclonal anti-Na/K ATPase (Santa Cruz, sc-48345; IB 1:1,000), rabbit monoclonal anti-Na/K ATPase (Abcam, ab-76020; IB 1:1,000; IF 1:400), mouse monoclonal anti-ERGIC53 (Novus, np62-03381; IF 1:400), mouse monoclonal anti-GM130 (BD biosciences, 610823; IF 1:400), sheep polyclonal anti-TGN46 (Bio-Rad, AHP1586; IF 1:400), mouse monoclonal anti-EEA1 (Cell Signaling, 48453; IF 1:400), mouse monoclonal anti-LAMP-1 (Santa Cruz, sc-20011; IB 1:1,000; IF 1:400), HRP-conjugated goat anti-rabbit IgG (Thermo Fisher Scientific, 31460; IB 1:5,000), HRP-conjugated goat anti-mouse IgG (Thermo Fisher Scientific, 31430; IB 1:5,000), HRP-conjugated donkey anti-goat IgG (Thermo Fisher Scientific; PA1-28664; IB 1:5,000), Dylight 488-conjugated donkey-anti-sheep/goat IgG (Bio-Rad, STAR88D488GA; IF 1:400), Cyanine Cy™2-conjugated donkey anti-mouse IgG (Jackson ImmunoResearch Laboratories, 715-225-150; IF 1:400); Cyanine Cy™2-conjugated donkey anti-rabbit IgG (Jackson ImmunoResearch Laboratories, 711-225-152; IF 1:400); Cyanine Cy™3-conjugated donkey anti-rabbit IgG (Jackson ImmunoResearch Laboratories, 715-165-152; IF 1:400); Cyanine Cy™3-conjugated donkey anti-mouse IgG (Jackson ImmunoResearch Laboratories, 715-165-150; IF 1:400); Cyanine Cy™3-conjugated donkey anti-goat IgG (Jackson ImmunoResearch Laboratories, 705-165-147; IF 1:400), Cyanine Cy™5-conjugated donkey anti-rabbit IgG (Jackson ImmunoResearch Laboratories, 711-175-152; IF 1:400) and Cyanine Cy™5-conjugated donkey anti-goat IgG (Jackson ImmunoResearch Laboratories, 705-175-147; IF 1:400).

### DNA constructs

pcDNA3.1(+) encoding *N*-terminal FLAG-tagged SMS2, SMS2^I62S^ and SMS2^M64R^ were described in (Pekkinen et al., 2019). DNA encoding *N*-terminal FLAG-tagged chimera SMSr-SMS211-77 was synthetically prepared (IDT, Belgium) and inserted into pcDNA3.1(+) using BamHI and NotI restriction sites. Pathogenic mutations were introduced using a QuickChangeII site-directed mutagenesis kit (Agilent Technologies, USA) and primers listed in Supplementary Table S1. To prepare lentiviral expression constructs, the ORF of FLAG-tagged SMS2 was PCR amplified using pcDNA3.1-FLAG-SMS2 as a template. The amplified DNA was inserted into pENTR™11 (Invitrogen, A10467) using the BamHI and NotI restriction sites. Pathogenic mutations and/or mutations affecting active site residue Asp276 were introduced by site-directed mutagenesis as described above. The inserts of the pENTR™11 constructs were transferred into the lentiviral expression vector pInducer20 (Addgene, 44012) using Gateway cloning (Invitrogen) according to the manufacturer’s instructions. The constructs encoding FLAG-tagged EqtSM (pET28a-EQ-SM-3xFLAG), GFP-tagged EqtSM_SS_ (pN1-EqtSM_SS_-oxGFP) and GFP-tagged EqtSol_SS_ (pN1-EqtSol_SS_-oxGFP) were described in (Deng et al., 2016). The constructs encoding GFP-tagged EqtSM_cyto_ (pN1-EqtSMcyto-oxGFP), GFP-tagged EqtSol_cyto_(pN1-EqtSol_cyto_-oxGFP) and mKate-tagged EqtSMcyto (pN1-EqtSMcyto-mkate) were described in (Niekamp *et al*., 2022). The construct encoding mCherry-tagged VAPA was described in (Jain et al., 2017). The construct encoding GFP-tagged Sec16L (pEFP-C1-Sec16L) was a kind gift from Benjamin Glick (University of Chicago, USA) and described in (Bhattacharyya and Glick, 2007). The expression construct encoding mCherry-tagged D4H (pN1-D4H-mCherry) was kindly provided by Gregory Fairn (University of Toronto, Canada) and described in (Maekawa and Fairn, 2015).

### Mammalian cell culture and transfection

Human fibroblasts derived from skin biopsies of OP-CDL patients and healthy controls were previously described in (Pekkinen et al., 2019). Human fibroblasts, human cervical carcinoma HeLa cells (ATCC CCL-2), human osteosarcoma epithelial U2OS cells (ATCC HTB-96), and human embryonic kidney 293 cells transformed with Simian Virus 40 large T antigen (HEK293T, ATCC CRL-3216) were cultured in high glucose Dulbecco’s modified Eagle’s medium (DMEM) containing 2 mM L-glutamine and 10% FBS, unless indicated otherwise. A HeLa cell-line lacking SMS1 and SMS2 (ΔSMS1/2) was described previously (Niekamp et al., 2022). DNA transfections were carried out using Lipofectamine 3000 (Thermo Fisher Scientific, L3000001) according to the manufacturer’s instructions.

### Lentiviral transduction

HeLa ΔSMS1/2 cells were stably transduced with pInducer20 constructs encoding FLAG-tagged SMS2, SMS2^D276A^, SMS2^I62S^, SMS2^I62S/D276A^, SMS2^M64R^ or SMS2^M64R/D276A^. To this end, low passage HEK293T cells were co-transfected with the corresponding pInducer20-FLAG-SMS2 construct and the packaging vectors psPAX2 (Addgene, 12260) and pMD2.G (Addgene, 12259). Culture medium was changed 6 h post-transfection. After 48 h, the lentivirus-containing medium was harvested, passed through a 0.45 μm filter, mixed 1:1 (v/v) with DMEM containing 8 μg/ml polybrene and used to infect HeLa ΔSMS1/2 cells. At 24 h post-infection, the medium was replaced with DMEM containing 1 mg/ml G418 and selective medium was changed daily. After 5 days, positively transduced cells were analyzed for doxycycline-dependent expression of the FLAG-tagged SMS2 variant using immunoblot analysis, immunofluorescence microscopy and metabolic labeling with clickSph, as described below.

### Cell lysis and immunoblot analysis

Cells were harvested and lysed in Lysis Buffer (1% TritonX-100, 1 mM EDTA pH 8.0, 150 mM NaCl, 20 mM Tris pH 7.5) supplemented with Protease Inhibitor Cocktail (PIC; 1 μg/ml aprotinin, 1 μg/ml leupeptin, 1 μg/ml pepstatin, 5 μg/ml antipain, 157 μg/ml benzamidine). Nuclei were removed by centrifugation at 600 x g for 10 min at 4°C. Post nuclear supernatants were collected and stored at – 80°C until use. Protein samples were mixed with 2× Laemmli Sample Buffer (0.3 M Tris HCl, pH 6.8, 10% SDS, 50% glycerol, 10% 2-β-mercaptoethanol, 0.025 % bromphenol blue), resolved by SDS-PAGE using 12% acrylamide gels, and transferred onto nitrocellulose membrane (0.45 μm; GE Health Sciences USA). Membranes were blocked with 5% non-fat milk solution for 40 min and washed with 0.05% Tween in PBS (PBST). Next, membrane was incubated for 2 h with primary antibody in PBST, washed three times with PBST and incubated with HRP-conjugated secondary antibody in PBST. After washing in PBST, the membranes were developed using enhanced chemiluminescence substrate (ECL; Thermo Fisher Scientific, USA). Images were recorded using a ChemiDoc XRS+ System (Bio-Rad, USA) and processed with Image Lab Software (BioRad, USA).

### Metabolic labelling and TLC analysis

Cells were metabolically labeled for 24 h with 4 μM clickable sphingosine in Opti-MEM reduced serum medium without Phenol red (Gibco, 11058). Next, cells were washed with PBS, harvested, and subjected to Bligh and Dyer lipid extraction (Bligh and Dyer, 1959). Dried lipid films were click reacted in a 40 μl reaction mix containing 0.45 mM fluorogenic dye 3-azido-7-hydroxycoumarin (Jena Bioscience, CLK-FA047), 1.4 mM Cu(I)tetra(acetonitrile) tetrafluoroborate and 66 % EtOH:CHCl_3_:CH_3_CN (66:19:16, v:v:v). Reaction mixtures were incubated at 40°C for 4 h followed by 12 h incubation at 12°C and applied at 120 nl/s to NANO-ADAMANT HP-TLC plates (Macherey-Nagel, Germany) with a CAMAG Linomat 5 TLC sampler (CAMAG, Switzerland). The TLC plate was developed in CHCl_3_:MeOH:H_2_O:AcOH 65:25:4:1, v:v:v:v) using a CAMAG ADC2 automatic TLC developer (CAMAG, Switzerland). The coumarin-derivatized lipids were visualized using a ChemiDoc XRS+ with UV-transillumination and Image Lab Software (BioRad, USA).

### Fluorescence microscopy

For immunofluorescence microscopy, cells were grown on glass coverslips and fixed in 4% paraformaldehyde (PFA) for 15 min at RT. After quenching in 50 mM ammonium chloride, cells were permeabilized with permeabilization buffer (PBS containing 0.3% (v/v) Triton-X100 and 1% (v/v) BSA) for 15 min. Immunostaining was performed in permeabilization buffer and nuclei were counterstained with DAPI, as described in (Jain et al., 2017). Coverslips were mounted onto glass slides using ProLong Gold Antifade Reagent (Thermo Fisher Scientific, USA). Fluorescence images were captured using a DeltaVision Elite Imaging System (GE Health Sciences, USA) or Leica DM5500B microscope (Leica, Germany), as indicated.

Imaging of live cells expressing SM or cholesterol reporters was performed using a Zeiss Cell Observer Spinning Disc Confocal Microscope equipped with a TempModule S1 temperature control unit, a Yokogawa Spinning Disc CSU-X1a 5000 Unit, a Evolve EMCDD camera (Photonics, Tucson), a motorized xyz-stage PZ-2000 XYZ (Applied Scientific Instrumentation) and an Alpha Plan-Apochromat x 63 (NA 1.46) oil immersion objective. The following filter combinations were used: blue emission with BP 445/50, green emission with BP 525/50, orange emission BP 605/70. All images were acquired using Zeiss Zen 2012 acquisition software. For hypotonic swelling, U2OS cells or fibroblasts were seeded in a μ-Slide 8 well glass bottom chamber (Ibidi; 80827) and transfected with indicated expression constructs. At 16h post-transfection, cells were imaged in isotonic medium (100% Opti-MEM) or after 5 min incubation in hypotonic medium (1% Opti-MEM in H_2_0) at 37°C. For cholesterol localization experiments, cells transfected with D4H-mCherry were incubated for 16 h in growth medium containing 70 μg/ml 10kDa dextran conjugated with Alexa Fluor 647 (Thermo Fisher Scientific, D22914). Growth medium was replaced with Opti-MEM 2 h prior to imaging. Images were deconvoluted using Huygens deconvolution (SVI, The Netherlands) and processed using Fiji software (NIH, USA).

Ratiometric confocal imaging of cells stained with Nile Red-based solvatochromic probes (NR12A, NRER_CI_) was performed on a Zeiss LSM 880 with an AiryScan module using a 63X 1.4 NA oil immersion objective. Excitation was provided by a 532 nm laser and fluorescence emission was detected at two spectral ranges: 500–600 (1500–600) and 600–700 nm (1600–700). The images were processed with a home-made program under LabView, which generates a ratiometric image by dividing the image of the 1500–600 channel by that of the 1600–700 channel, as described in (Darwich et al., 2013). For each pixel, a pseudo-color scale was used for coding the ratio, while the intensity was defined by the integral intensity recorded for both channels at the corresponding pixel. For staining with the NR12A probe, cells were washed with PBS and incubated in Opti-MEM containing 0.2 μM NR12A for 10 min at RT. For staining with the NRER_CI_ probe, cells were washed with PBS and incubated in Opti-MEM containing 0.2 μM NRER_CI_ for 30 min at 37°C. Subsequently, cells were washed 3 times with PBS and imaged in Opti-MEM.

### Cytotoxicity assay

Cells were seeded in 96-well plate (Greiner Bio-One; 655101) at 10,000 cells per well in DMEM supplemented with 10% FBS. After 24 h, the medium was replaced with Opti-MEM, and 24 h later mβCD was added at the indicated concentrations. After 1 h, PrestoBlue HS (Thermo Fisher Scientific; P50200) was added to the well to a final concentration of 10% (v/v) and incubated for 3.5 h at 37°C. Next, absorbance at 570 nm was measured with 600 nm as reference wavelength using an Infinite 200 Pro M-Plex plate reader (Tecan Lifesciences). Measurements were average of quadruplicates. To calculate relative percentage of cell survival, the measured value for each well (x) was subtracted by the minimum measured value (min) and divided by the subtrahend of the average measured value of untreated cells (untreated) and the minimum measured value (min); ((x-min)/(untreated-min) *100).

### Cell surface staining with EqtSM

For production of recombinant FLAG-tagged EqtSM, *E. coli* BL21 (DE3) transformed with pET28a-EQ-SM-3xFLAG was grown at 37°C to early exponential phase in LB medium containing 100 μg/ml ampicillin prior to addition of 0.4 mM isopropyl β-D-1-thiogalactopyranoside. After 5 h induction, bacteria were mechanically lysed in 20 mM Na_2_HPO_4_/NaH_2_PO_4_, pH 7.4, 500 mM NaCl, and 25 mM imidazole supplemented with PIC by microtip sonication. Bacterial lysates were cleared by centrifugation at 10,000 *g* for 20 min at 4°C and applied to a HisTrap HP column using an AKTA Prime protein purification system (GE Healthcare, Life Sciences). Bound protein was eluted with a linear imidazole gradient. HeLa cells grown on glass coverslips were incubated with 1 μM of purified FLAG-EqtSM in Labeling Buffer (20 mM Na_2_HPO_4_/NaH_2_PO_4_, pH 7.4, 500 mM NaCl) for 2 min at RT, washed with PBS, and then fixed in 4% PFA for 15 min at RT. After quenching in 50 mM ammonium chloride, cells were immunostained with polyclonal rabbit anti-FLAG antibody at 4°C for 10 min and Cy2-conjugated anti-rabbit antibody at RT for 30 min in PBS supplemented with 1% BSA. For flow cytometry, HeLa cells were trypsinized, resuspended in medium containing 10% FCS, washed and then incubated with 1 μM Eqt-FLAG at RT for 2 min in Labeling Buffer. Next, cells were incubated with anti-FLAG antibody for 10 min at 4°C in PBS containing 1% BSA, washed, and then fixed in 4% PFA for 15 min at RT. After quenching in 50 mM ammonium chloride, cells were incubated with Cy2-conjugated anti-rabbit antibody for 45 min in PBS containing 1% BSA, washed, and subjected to flow cytometry using a SH800 Cell Sorter (Sony Biotechnology). Flow cytometry data were analysed using Sony Cell Sorter software version 2.1.5.

### LC-MS/MS lipidomics

Cells were incubated for 24h in Opti-MEM reduced serum medium in the absence or presence of 1μg/ml doxycycline. Next, cells were harvested in homogenization buffer (15 mM KCl, 5 mM NaCl, 20 mM HEPES/KOH pH 7.2, 10% glycerol, 1x PIC) using a sonifier BRANSON 250.The protein in crude homogenates was determined by Bradford protein assay (BioRad, USA) and 50 μg of protein was used for a subsequent chloroform/methanol extraction. To normalize lipid concentration of lipids in the samples, homogenates were prior to the extraction spiked with lipid standards ceramide (Cer 18:1/17:0) and sphingomyelin (SM 18:1/17:0). Dried lipid extracts were dissolved in a 50:50 mixture of mobile phase A (60:40 water/acetonitrile, including 10 mM ammonium formate and 0.1% formic acid) and mobile phase B (88:10:2 2-propanol/acetonitrile/H_2_0, including 2 mM ammonium formate and 0.02% formic acid). HPLC analysis was performed on a C30 reverse-phase column (Thermo Acclaim C30, 2.1 × 250 mm, 3 μm, operated at 50°C; Thermo Fisher Scientific) connected to an HP 1100 series HPLC system(Agilent) and a *QExactivePLUS* Orbitrap mass spectrometer (Thermo Fisher Scientific) equipped with a heated electrospray ionization (HESI) probe. MS analysis was performed as described previously (Eising et al., 2019). Briefly, elution was performed with a gradient of 45 min; during the first 3 min, elution started with 40% of phase B and increased to 100% in a linear gradient over 23 mins. 100% of B was maintained for 3 min. Afterwards, solvent B was decreased to 40% and maintained for another 15 min for column re-equilibration. MS spectra of lipids were acquired in full-scan/data-dependent MS2 mode. The maximum injection time for full scans was 100 ms, with a target value of 3,000,000 at a resolution of 70,000 at m/z 200 and a mass range of 200–2000 m/z in both positive and negative mode. The 10 most intense ions from the survey scan were selected and fragmented with HCD with a normalized collision energy of 30. Target values for MS/MS were set at 100,000 with a maximum injection time of 50 ms at a resolution of 17,500 at m/z of 200. Peaks were analyzed using the Lipid Search algorithm (MKI, Tokyo, Japan). Peaks were defined through raw files, product ion and precursor ion accurate masses. Candidate molecular species were identified by database (>1,000,000 entries) search of positive (+H^+^; +NH_4_) or negative ion adducts (–H^−^; +COOH^−^). Mass tolerance was set to 5 ppm for the precursor mass. Samples were aligned within a time window and results combined in a single report. From the intensities of lipid standards and lipid classes used, absolute values for each lipid in pmol/mg protein were calculated. Data are reported as mol% of total phospholipids measured.

### Organellar purification and shot-gun lipid mass spectrometry

#### Affinity purification

10 million cells were seeded in 15 cm dishes and cultured for 24 h in Opti-MEM reduced serum medium containing 1 μg/ml doxycycline. For ER purifications, cells were washed once in ice-cold PBS. For PM purification, cells were washed three times with ice-cold PBS and then incubated for 30 min at 4°C in PBS containing 1 mg/ml EZ-Link-sulfo-NHS-LC-LC-Biotin (Thermo Fisher Scientific). Next, cells were washed three times with ice-cold PBS. Biotin-treated and untreated cells were scraped in ice-cold PBS, centrifuged once at 500 × g, and then twice at 1,000 × g for 5 min, 4°C. All steps from here were performed on ice or at 4°C. Cell pellets were re-suspended in ice-cold 10 ml SuMa buffer [10 mM Hepes, 0.21 M mannitol, 0.070 M Sucrose, pH 7.5]. After the third centrifugation step, cells were resuspended in 1 ml SuMa4 buffer [SuMa buffer supplemented with 0.5 mM DTT, 0.5 % fatty acid-free BSA (Sigma Aldrich), 25 units/ml Benzonase (Sigma Aldrich) and 1x cOmplete™ Mini, EDTA-free Protease Inhibitor Cocktail (Roche Diagnostics) and lysed by passages through a Balch homogenizer 20 times. The cell lysates were subsequently centrifuged at 1,500 × g for 10 min and again for 15 min to prepare the light membrane fractions (LMFs). For ER purification, LMFs were incubated with rabbit anti-calnexin polyclonal antibody (Abcam) and subsequently with anti-rabbit IgG MicroBeads (Miltenyi Biotec), and for the PM purification, only with Streptavidin MicroBeads (Miltenyi Biotec). The LMFs were then loaded into MS Columns (Miltenyi Biotec) mounted on a magnetic stand (Miltenyi Biotec) and pre-equilibrated in SuMa4 buffer. The columns were washed three times with 500 μl SuMa4 buffer and twice with 500 μl SuMa2 buffer (SuMa4 without Benzonase and PIC). Thereafter, columns were removed from the magnetic stand and elution was performed with 600 μl SuMa2 buffer. Eluate samples were equally divided for western blotting and lipidomics. All samples were centrifuged at 21,100 × g for 20 min and supernatants were discarded. Pellets were resuspended in 200 μl SuMa+ buffer (SuMa2 without BSA) and the centrifugation was repeated. Lipidomics samples were stored at −80°C until analysis and immunoblot samples were dissolved in 2× Laemmli Sample Buffer containing 100 mM DTT and stored at −20 °C until processing.

#### Lipid extraction

Lipid extraction was performed as previously described (Nielsen et al., 2020), with some modifications. Briefly, samples in 200 μl 155 mM ammonium bicarbonate were mixed with 24 μl internal lipid standard mix (Nielsen et al., 2020) and 976 μl chloroform:methanol 2:1 (*v/v*). The samples were shaken in a thermomixer at 2,000 rpm and 4°C for 15 min and centrifuged for 2 min at 2,000 × g and 4°C. Then, the lower phase containing lipids was washed twice with 100 μl methanol and 50 μl 155 mM ammonium bicarbonate. Lower phase was then transferred to new tubes and dried in a vacuum centrifuge for 75 min, and the dried lipids were resuspended in 100 μl chloroform:methanol 1:2 (*v:v*).

#### Mass spectrometry

Shotgun lipidomics was performed as previously described (Nielsen et al., 2020). Lipid extracts (10 μl) were loaded in a 96 well plate and mixed with either 12.9 μl positive ionization solvent (13.3 mM ammonium acetate in propan-2-ol) or 10 μl negative ionization solvent (0.2 % (*v/v*) triethyl amine in chloroform:methanol 1:5 (*v/v*)). The samples were analyzed in the negative and positive ionization modes using Q Exactive Hybrid Quadrupole-Orbitrap mass spectrometer (Thermo Fisher Scientific, Waltham, MA) coupled to TriVersa NanoMate (Advion Biosciences, Ithaca, NY, USA). Data are reported as mol% of total lipids measured.

## ACKNOWLEDGEMENTS

We gratefully acknowledge Gregory Fairn, Benjamin Glick and Christoph Thiele for providing DNA constructs and chemicals, Florian Fröhlich and Stefan Walter for technical assistance with LC-MS/MS, and Rainer Kurre for technical assistance with live cell microscopy. This work was supported by the Deutsche Forschungsgemeinschaft (SFB944-P14 and HO3539/1-1 to J.C.M.H.; SFB944-P8 to J.Pi.) and the National Institute of General Medical Sciences of the United States National Institutes of Health (award R35 GM144096 to C. G. B.). The authors declare no competing interests.

## AUTHOR CONTRIBUTIONS

T.S., J.Pa. and J.C.M.H. designed the research plan and wrote the manuscript; T.S. and J.Pa. performed the bulk of experiments and analyzed the results, with critical input from M.B. and A.H.; D.I.D. and A.S.K. provided Nile Red-based solvatochromic probes and critical expertise on their use; B.S. performed the experiments with Nile Red-based solvatochromic probes, with critical input from T.S., M.P. and J.Pi.; L.D.B. and P.A.T. provided patient-derived fibroblasts; O.M. provided patient-relevant expertise and helped to interpret experimental data; Y.K. and C.G.B. designed and characterized the Eqt-based SM reporters; M.M.F. and K.M. performed shotgun lipidomics on isolated organelles; all authors discussed results and commented on the manuscript.

## REFERENCES

Alonso, A., and F.M. Goñi. 2018. The Physical Properties of Ceramides in Membranes. Annu. Rev. Biophys. 47:633–654. doi:10.1146/annurev-biophys-070317-033309.

Aubin, I., C.P. Adams, S. Opsahl, D. Septier, C.E. Bishop, N. Auge, R. Salvayre, A. Negre-Salvayre, M. Goldberg, J.-L. Guénet, and C. Poirier. 2005. A deletion in the gene encoding sphingomyelin phosphodiesterase 3 (Smpd3) results in osteogenesis and dentinogenesis imperfecta in the mouse. Nat. Genet. 37:803–5. doi:10.1038/ng1603.

Bhattacharyya, D., and B.S. Glick. 2007. Two Mammalian Sec16 Homologues Have Nonredundant Functions in Endoplasmic Reticulum (ER) Export and Transitional ER Organization. Mol. Biol. Cell. 18:839–849. doi:10.1091/mbc.e06-08-0707.

Bigay, J., and B. Antonny. 2012. Curvature, lipid packing, and electrostatics of membrane organelles: defining cellular territories in determining specificity. Dev. Cell. 23:886–95. doi:10.1016/j.devcel.2012.10.009.

Boyadjiev, S.A., J.C. Fromme, J. Ben, S.S. Chong, C. Nauta, D.J. Hur, G. Zhang, S. Hamamoto, R. Schekman, M. Ravazzola, L. Orci, and W. Eyaid. 2006. Cranio-lenticulo-sutural dysplasia is caused by a SEC23A mutation leading to abnormal endoplasmic-reticulum-to-Golgi trafficking. Nat. Genet. 38:1192–7. doi:10.1038/ng1876.

Brügger, B., R. Sandhoff, S. Wegehingel, K. Gorgas, J. Malsam, J.B. Helms, W.D. Lehmann, W. Nickel, and F.T. Wieland. 2000. Evidence for segregation of sphingomyelin and cholesterol during formation of COPI-coated vesicles. J. Cell Biol. 151:507–18. doi:10.1083/jcb.151.3.507.

Buton, X., P. Hervé, J. Kubelt, A. Tannert, K.N.J. Burger, P. Fellmann, P. Müller, A. Herrmann, M. Seigneuret, and P.F. Devaux. 2002. Transbilayer movement of monohexosylsphingolipids in endoplasmic reticulum and Golgi membranes. Biochemistry. 41:13106–15. doi:10.1021/bi020385t.

Chalat, M., I. Menon, Z. Turan, and A.K. Menon. 2012. Reconstitution of glucosylceramide flip-flop across endoplasmic reticulum: implications for mechanism of glycosphingolipid biosynthesis. J. Biol. Chem. 287:15523–32. doi:10.1074/jbc.M112.343038.

Chung, J., F. Torta, K. Masai, L. Lucast, H. Czapla, L.B. Tanner, P. Narayanaswamy, M.R. Wenk, F. Nakatsu, and P. De Camilli. 2015. INTRACELLULAR TRANSPORT. PI4P/phosphatidylserine countertransport at ORP5- and ORP8-mediated ER-plasma membrane contacts. Science. 349:428–32. doi:10.1126/science.aab1370.

Danylchuk, D.I., P.-H. Jouard, and A.S. Klymchenko. 2021. Targeted Solvatochromic Fluorescent Probes for Imaging Lipid Order in Organelles under Oxidative and Mechanical Stress. J. Am. Chem. Soc. 143:912–924. doi:10.1021/jacs.0c10972.

Danylchuk, D.I., S. Moon, K. Xu, and A.S. Klymchenko. 2019. Switchable Solvatochromic Probes for Live-Cell Super-resolution Imaging of Plasma Membrane Organization. Angew. Chem. Int. Ed. Engl. 58:14920–14924. doi:10.1002/anie.201907690.

Darwich, Z., O.A. Kucherak, R. Kreder, L. Richert, R. Vauchelles, Y. Mély, and A.S. Klymchenko. 2013. Rational design of fluorescent membrane probes for apoptosis based on 3-hydroxyflavone. Methods Appl. Fluoresc. 1:025002. doi:10.1088/2050-6120/1/2/025002.

Das, A., M.S. Brown, D.D. Anderson, J.L. Goldstein, and A. Radhakrishnan. 2014. Three pools of plasma membrane cholesterol and their relation to cholesterol homeostasis. Elife. 3. doi:10.7554/eLife.02882.

Deng, Y., F.E. Rivera-Molina, D.K. Toomre, and C.G. Burd. 2016. Sphingomyelin is sorted at the trans Golgi network into a distinct class of secretory vesicle. Proc. Natl. Acad. Sci. U. S. A. 113:6677–82. doi:10.1073/pnas.1602875113.

Ehlen, H.W.A., M. Chinenkova, M. Moser, H.-M. Munter, Y. Krause, S. Gross, B. Brachvogel, M. Wuelling, U. Kornak, and A. Vortkamp. 2013. Inactivation of anoctamin-6/Tmem16f, a regulator of phosphatidylserine scrambling in osteoblasts, leads to decreased mineral deposition in skeletal tissues. J. Bone Miner. Res. 28:246–59. doi:10.1002/jbmr.1751.

Eising, S., L. Thiele, and F. Fröhlich. 2019. A systematic approach to identify recycling endocytic cargo depending on the GARP complex. Elife. 8. doi:10.7554/eLife.42837.

Ellison, C.J., W. Kukulski, K.B. Boyle, S. Munro, and F. Randow. 2020. Transbilayer Movement of Sphingomyelin Precedes Catastrophic Breakage of Enterobacteria-Containing Vacuoles. Curr. Biol. 30:2974–2983.e6. doi:10.1016/j.cub.2020.05.083.

Endapally, S., D. Frias, M. Grzemska, A. Gay, D.R. Tomchick, and A. Radhakrishnan. 2019. Molecular Discrimination between Two Conformations of Sphingomyelin in Plasma Membranes. Cell. 176:1040–1053.e17. doi:10.1016/j.cell.2018.12.042.

Fukasawa, M., M. Nishijima, H. Itabe, T. Takano, and K. Hanada. 2000. Reduction of sphingomyelin level without accumulation of ceramide in Chinese hamster ovary cells affects detergent-resistant membrane domains and enhances cellular cholesterol efflux to methyl-beta - cyclodextrin. J. Biol. Chem. 275:34028–34. doi:10.1074/jbc.M005151200.

Garbes, L., K. Kim, A. Rieß, H. Hoyer-Kuhn, F. Beleggia, A. Bevot, M.J. Kim, Y.H. Huh, H.-S. Kweon, R. Savarirayan, D. Amor, P.M. Kakadia, T. Lindig, K.O. Kagan, J. Becker, S.A. Boyadjiev, B. Wollnik, O. Semler, S.K. Bohlander, J. Kim, and C. Netzer. 2015. Mutations in SEC24D, encoding a component of the COPII machinery, cause a syndromic form of osteogenesis imperfecta. Am. J. Hum. Genet. 96:432–9. doi:10.1016/j.ajhg.2015.01.002.

Ghanbarpour, A., D.P. Valverde, T.J. Melia, and K.M. Reinisch. 2021. A model for a partnership of lipid transfer proteins and scramblases in membrane expansion and organelle biogenesis. Proc. Natl. Acad. Sci. U. S. A. 118. doi:10.1073/pnas.2101562118.

Granado, M.H., P. Gangoiti, A. Ouro, L. Arana, and A. Gómez-Muñoz. 2009. Ceramide 1-phosphate inhibits serine palmitoyltransferase and blocks apoptosis in alveolar macrophages. Biochim. Biophys. Acta. 1791:263–72. doi:10.1016/j.bbalip.2009.01.023.

Guillemyn, B., S. Nampoothiri, D. Syx, F. Malfait, and S. Symoens. 2021. Loss of TANGO1 Leads to Absence of Bone Mineralization. JBMR plus. 5:e10451. doi:10.1002/jbm4.10451.

Hanada, K., K. Kumagai, S. Yasuda, Y. Miura, M. Kawano, M. Fukasawa, and M. Nishijima. 2003. Molecular machinery for non-vesicular trafficking of ceramide. Nature. 426:803–9. doi:10.1038/nature02188.

Holthuis, J.C.M., and A.K. Menon. 2014. Lipid landscapes and pipelines in membrane homeostasis. Nature. 510. doi:10.1038/nature13474.

Huang, D., B. Xu, L. Liu, L. Wu, Y. Zhu, A. Ghanbarpour, Y. Wang, F.-J. Chen, J. Lyu, Y. Hu, Y. Kang, W. Zhou, X. Wang, W. Ding, X. Li, Z. Jiang, J. Chen, X. Zhang, H. Zhou, J.Z. Li, C. Guo, W. Zheng, X. Zhang, P. Li, T. Melia, K. Reinisch, and X.-W. Chen. 2021. TMEM41B acts as an ER scramblase required for lipoprotein biogenesis and lipid homeostasis. Cell Metab. 33:1655–1670.e8. doi:10.1016/j.cmet.2021.05.006.

Huitema, K., J. Van Den Dikkenberg, J.F.H.M. Brouwers, and J.C.M. Holthuis. 2004. Identification of a family of animal sphingomyelin synthases. EMBO J. 23. doi:10.1038/sj.emboj.7600034.

Jain, A., O. Beutel, K. Ebell, S. Korneev, and J.C.M. Holthuis. 2017. Diverting CERT-mediated ceramide transport to mitochondria triggers Bax-dependent apoptosis. J. Cell Sci. 130. doi:10.1242/jcs.194191.

Kim, Y.-J., P. Greimel, and Y. Hirabayashi. 2018. GPRC5B-Mediated Sphingomyelin Synthase 2 Phosphorylation Plays a Critical Role in Insulin Resistance. iScience. 8:250–266. doi:10.1016/j.isci.2018.10.001.

King, C., P. Sengupta, A.Y. Seo, and J. Lippincott-Schwartz. 2020. ER membranes exhibit phase behavior at sites of organelle contact. Proc. Natl. Acad. Sci. U. S. A. 117:7225–7235. doi:10.1073/pnas.1910854117.

Lamour, N.F., P. Subramanian, D.S. Wijesinghe, R. V Stahelin, J. V Bonventre, and C.E. Chalfant. 2009. Ceramide 1-phosphate is required for the translocation of group IVA cytosolic phospholipase A2 and prostaglandin synthesis. J. Biol. Chem. 284:26897–907. doi:10.1074/jbc.M109.001677.

Levental, K.R., E. Malmberg, J.L. Symons, Y.-Y. Fan, R.S. Chapkin, R. Ernst, and I. Levental. 2020. Lipidomic and biophysical homeostasis of mammalian membranes counteracts dietary lipid perturbations to maintain cellular fitness. Nat. Commun. 11:1339. doi:10.1038/s41467-020-15203-1.

Li, Z., H. Zhang, J. Liu, C.-P. Liang, Y. Li, Y. Li, G. Teitelman, T. Beyer, H.H. Bui, D.A. Peake, Y. Zhang, P.E. Sanders, M.-S. Kuo, T.-S. Park, G. Cao, and X.-C. Jiang. 2011. Reducing plasma membrane sphingomyelin increases insulin sensitivity. Mol. Cell. Biol. 31:4205–18. doi:10.1128/MCB.05893-11.

Maekawa, M., and G.D. Fairn. 2015. Complementary probes reveal that phosphatidylserine is required for the proper transbilayer distribution of cholesterol. J. Cell Sci. 128:1422–33. doi:10.1242/jcs.164715.

Magdeleine, M., R. Gautier, P. Gounon, H. Barelli, S. Vanni, and B. Antonny. 2016. A filter at the entrance of the Golgi that selects vesicles according to size and bulk lipid composition. Elife. 5. doi:10.7554/eLife.16988.

van Meer, G., D.R. Voelker, and G.W. Feigenson. 2008. Membrane lipids: where they are and how they behave. Nat. Rev. Mol. Cell Biol. 9:112–24. doi:10.1038/nrm2330.

Mesmin, B., J. Bigay, J. Moser von Filseck, S. Lacas-Gervais, G. Drin, and B. Antonny. 2013. A four-step cycle driven by PI(4)P hydrolysis directs sterol/PI(4)P exchange by the ER-Golgi tether OSBP. Cell. 155:830–43. doi:10.1016/j.cell.2013.09.056.

Mesmin, B., J. Bigay, J. Polidori, D. Jamecna, S. Lacas-Gervais, and B. Antonny. 2017. Sterol transfer, PI4P consumption, and control of membrane lipid order by endogenous OSBP. EMBO J. 36:3156–3174. doi:10.15252/embj.201796687.

Mitsutake, S., K. Zama, H. Yokota, T. Yoshida, M. Tanaka, M. Mitsui, M. Ikawa, M. Okabe, Y. Tanaka, T. Yamashita, H. Takemoto, T. Okazaki, K. Watanabe, and Y. Igarashi. 2011. Dynamic modification of sphingomyelin in lipid microdomains controls development of obesity, fatty liver, and type 2 diabetes. J. Biol. Chem. 286:28544–55. doi:10.1074/jbc.M111.255646.

Moser von Filseck, J., A. Čopič, V. Delfosse, S. Vanni, C.L. Jackson, W. Bourguet, and G. Drin. 2015. INTRACELLULAR TRANSPORT. Phosphatidylserine transport by ORP/Osh proteins is driven by phosphatidylinositol 4-phosphate. Science. 349:432–6. doi:10.1126/science.aab1346.

Munro, S. 1995. An investigation of the role of transmembrane domains in Golgi protein retention. EMBO J. 14:4695–704.

Murakami, C., and F. Sakane. 2021. Sphingomyelin synthase-related protein generates diacylglycerol via the hydrolysis of glycerophospholipids in the absence of ceramide. J. Biol. Chem. 296:100454. doi:10.1016/j.jbc.2021.100454.

Murshed, M. 2018. Mechanism of Bone Mineralization. Cold Spring Harb. Perspect. Med. 8. doi:10.1101/cshperspect.a031229.

Niekamp, P., F. Scharte, T. Sokoya, L. Vittadello, Y. Kim, Y. Deng, E. Südhoff, A. Hilderink, M. Imlau, C. J. Clarke, M. Hensel, C. G. Burd, and J. C. M. Holthuis. 2022. Ca2+-activated sphingomyelin scrambling and turnover mediate ESCRT-independent lysosomal repair. Nature Commun. In press. doi:10.1038/s41467-022-29481-4.

Nielsen, I.Ø., A. Vidas Olsen, J. Dicroce-Giacobini, E. Papaleo, K.K. Andersen, M. Jäättelä, K. Maeda, and M. Bilgin. 2020. Comprehensive Evaluation of a Quantitative Shotgun Lipidomics Platform for Mammalian Sample Analysis on a High-Resolution Mass Spectrometer. J. Am. Soc. Mass Spectrom. 31:894–907. doi:10.1021/jasms.9b00136.

Nilsson, I., H. Ohvo-Rekilä, J.P. Slotte, A.E. Johnson, and G. von Heijne. 2001. Inhibition of protein translocation across the endoplasmic reticulum membrane by sterols. J. Biol. Chem. 276:41748–54. doi:10.1074/jbc.M105823200.

Di Paolo, G., and P. De Camilli. 2006. Phosphoinositides in cell regulation and membrane dynamics. Nature. 443:651–7. doi:10.1038/nature05185.

Pekkinen, M., P.A. Terhal, L.D. Botto, P. Henning, R.E. Mäkitie, P. Roschger, A. Jain, M. Kol, M.A. Kjellberg, E.P. Paschalis, K. van Gassen, M. Murray, P. Bayrak-Toydemir, M.K. Magnusson, J. Jans, M. Kausar, J.C. Carey, P. Somerharju, U.H. Lerner, V.M. Olkkonen, K. Klaushofer, J.C.M. Holthuis, and O. Mäkitie. 2019. Osteoporosis and skeletal dysplasia caused by pathogenic variants in SGMS2. JCI Insight. doi:10.1172/jci.insight.126180.

Pomorski, T.G., and A.K. Menon. 2016. Lipid somersaults: Uncovering the mechanisms of protein-mediated lipid flipping. Prog. Lipid Res. 64:69–84. doi:10.1016/j.plipres.2016.08.003.

Presa, N., A. Gomez-Larrauri, A. Dominguez-Herrera, M. Trueba, and A. Gomez-Muñoz. 2020. Novel signaling aspects of ceramide 1-phosphate. Biochim. Biophys. acta. Mol. cell Biol. lipids. 1865:158630. doi:10.1016/j.bbalip.2020.158630.

Quiroga, R., A. Trenchi, A. González Montoro, J. Valdez Taubas, and H.J.F. Maccioni. 2013. Short transmembrane domains with high-volume exoplasmic halves determine retention of Type II membrane proteins in the Golgi complex. J. Cell Sci. 126:5344–9. doi:10.1242/jcs.130658.

Radanović, T., J. Reinhard, S. Ballweg, K. Pesek, and R. Ernst. 2018. An Emerging Group of Membrane Property Sensors Controls the Physical State of Organellar Membranes to Maintain Their Identity. Bioessays. 40:e1700250. doi:10.1002/bies.201700250.

Roux, A., D. Cuvelier, P. Nassoy, J. Prost, P. Bassereau, and B. Goud. 2005. Role of curvature and phase transition in lipid sorting and fission of membrane tubules. EMBO J. 24:1537–45. doi:10.1038/sj.emboj.7600631.

Sharpe, H.J., T.J. Stevens, and S. Munro. 2010. A comprehensive comparison of transmembrane domains reveals organelle-specific properties. Cell. 142:158–69. doi:10.1016/j.cell.2010.05.037.

Slotte, J.P. 2013. Biological functions of sphingomyelins. Prog. Lipid Res. 52:424–37. doi:10.1016/j.plipres.2013.05.001.

Spandl, J., D.J. White, J. Peychl, and C. Thiele. 2009. Live cell multicolor imaging of lipid droplets with a new dye, LD540. Traffic. 10:1579–84. doi:10.1111/j.1600-0854.2009.00980.x.

Sugimoto, M., Y. Shimizu, S. Zhao, N. Ukon, K. Nishijima, M. Wakabayashi, T. Yoshioka, K. Higashino, Y. Numata, T. Okuda, N. Tamaki, H. Hanamatsu, Y. Igarashi, and Y. Kuge. 2016. Characterization of the role of sphingomyelin synthase 2 in glucose metabolism in whole-body and peripheral tissues in mice. Biochim. Biophys. Acta. 1861:688–702. doi:10.1016/j.bbalip.2016.04.019.

Vacaru, A.M., F.G. Tafesse, P. Ternes, V. Kondylis, M. Hermansson, J.F.H.M. Brouwers, P. Somerharju, C. Rabouille, and J.C.M. Holthuis. 2009. Sphingomyelin synthase-related protein SMSr controls ceramide homeostasis in the ER. J. Cell Biol. 185:1013–1027. doi:10.1083/jcb.200903152.

Wong, L.H., A.T. Gatta, and T.P. Levine. 2019. Lipid transfer proteins: the lipid commute via shuttles, bridges and tubes. Nat. Rev. Mol. Cell Biol. 20:85–101. doi:10.1038/s41580-018-0071-5.

Yano, M., K. Watanabe, T. Yamamoto, K. Ikeda, T. Senokuchi, M. Lu, T. Kadomatsu, H. Tsukano, M. Ikawa, M. Okabe, S. Yamaoka, T. Okazaki, H. Umehara, T. Gotoh, W.-J. Song, K. Node, R. Taguchi, K. Yamagata, and Y. Oike. 2011. Mitochondrial dysfunction and increased reactive oxygen species impair insulin secretion in sphingomyelin synthase 1-null mice. J. Biol. Chem. 286:3992–4002. doi:10.1074/jbc.M110.179176.

Yano, M., T. Yamamoto, N. Nishimura, T. Gotoh, K. Watanabe, K. Ikeda, Y. Garan, R. Taguchi, K. Node, T. Okazaki, and Y. Oike. 2013. Increased oxidative stress impairs adipose tissue function in sphingomyelin synthase 1 null mice. PLoS One. 8:e61380. doi:10.1371/journal.pone.0061380.

Zhou, Y., and J.F. Hancock. 2018. Deciphering lipid codes: K-Ras as a paradigm. Traffic. 19:157–165. doi:10.1111/tra.12541.

