## Supplementary material for "Pathogenic variants of sphingomyelin synthase SMS2 disrupt lipid landscapes in the secretory pathway": Figures S1-5; Table S1

**This PDF file includes:**

Supplementary Table 1

Supplementary Figures S1 to S5

**Supplementary Table 1. Primers used for site-directed mutagenesis of SMS2.**

| Primer name | Sequence |
|---|---|
| SMS2(I62S)-F | 5'-CGGACTATATCCAAAGTGCTATGCCCACTGAATC-3' |
| SMS2(I62S)-R | 5'-GATTCAGTGGGCATAGCACTTTGGATATAGTCCG-3' |
| SMS2(M64R)-F | 5'-CTATATCCAAATTGCTAGGCCCACTGAATCAAGG-3' |
| SMS2(M64R)-R | 5'-CCTTGATTCAGTGGGCCTAGCAATTTGGATATAG-3' |
| SMS2(D276A)-F | 5'-CGAACACTACACTATCGCTGTGATCATTGC-3' |
| SMS2(D276A)-R | 5'-GCAATGATCACAGCGATAGTGTAGTGTTCG-3' |

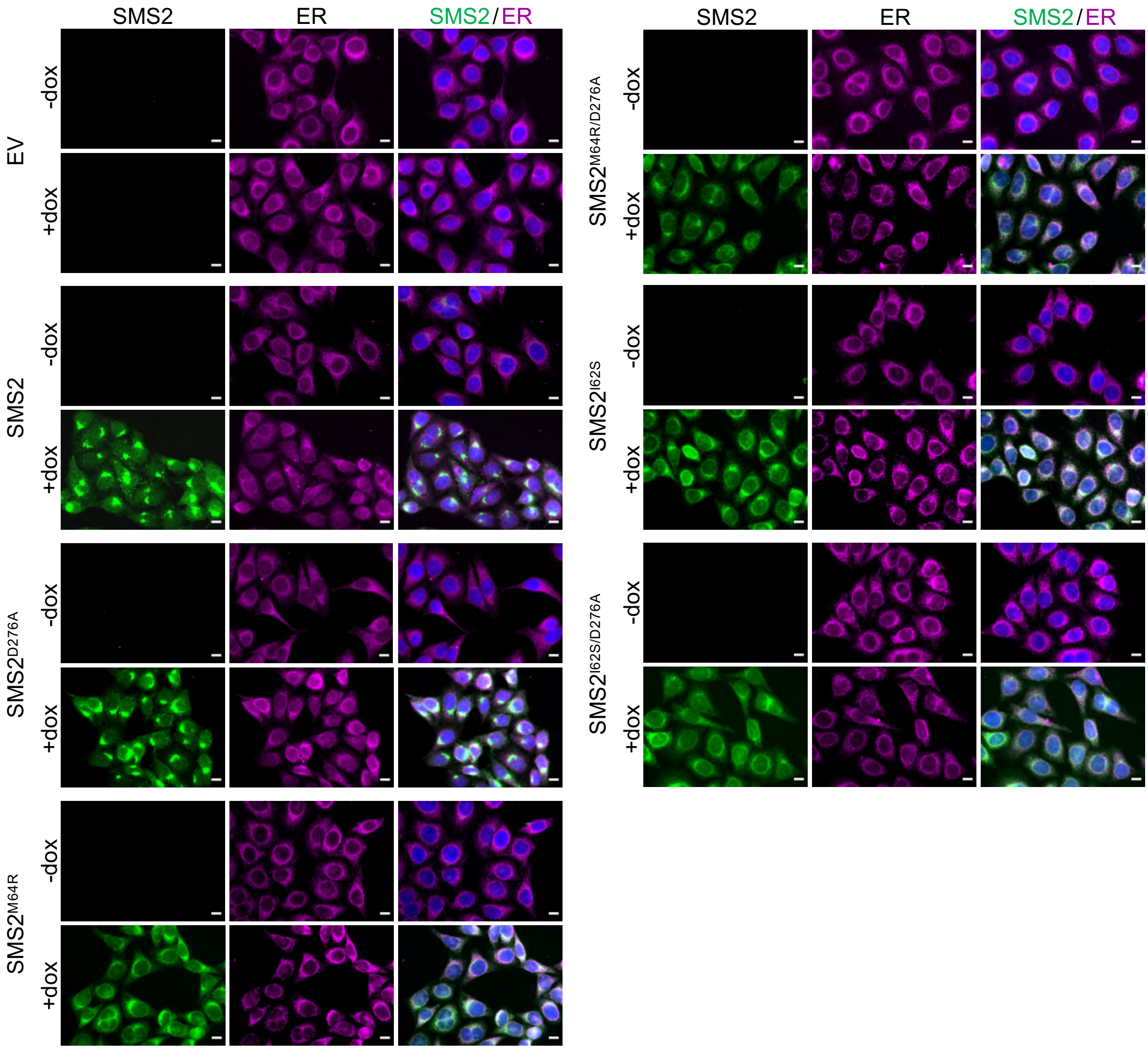


**Supplementary Figure 1. Doxycycline-induced expression of SMS2 variants in stably transduced HeLa cells.**

HeLa ΔSMS1/2 cells transduced with doxycycline-inducible Flag-tagged SMS2, $SMS2^{I62S}$, $SMS2^{M64R}$ or their enzyme-dead isoforms (D276A) were grown for 16 h in the absence or presence of 1 µg/ml doxycycline. Next, cells were fixed, immunostained with α-FLAG (*green*) and α-calnexin (*magenta*) antibodies, counterstained with DAPI (*blue*) and imaged by conventional fluorescence microscopy. Scale bar, 10 µm.

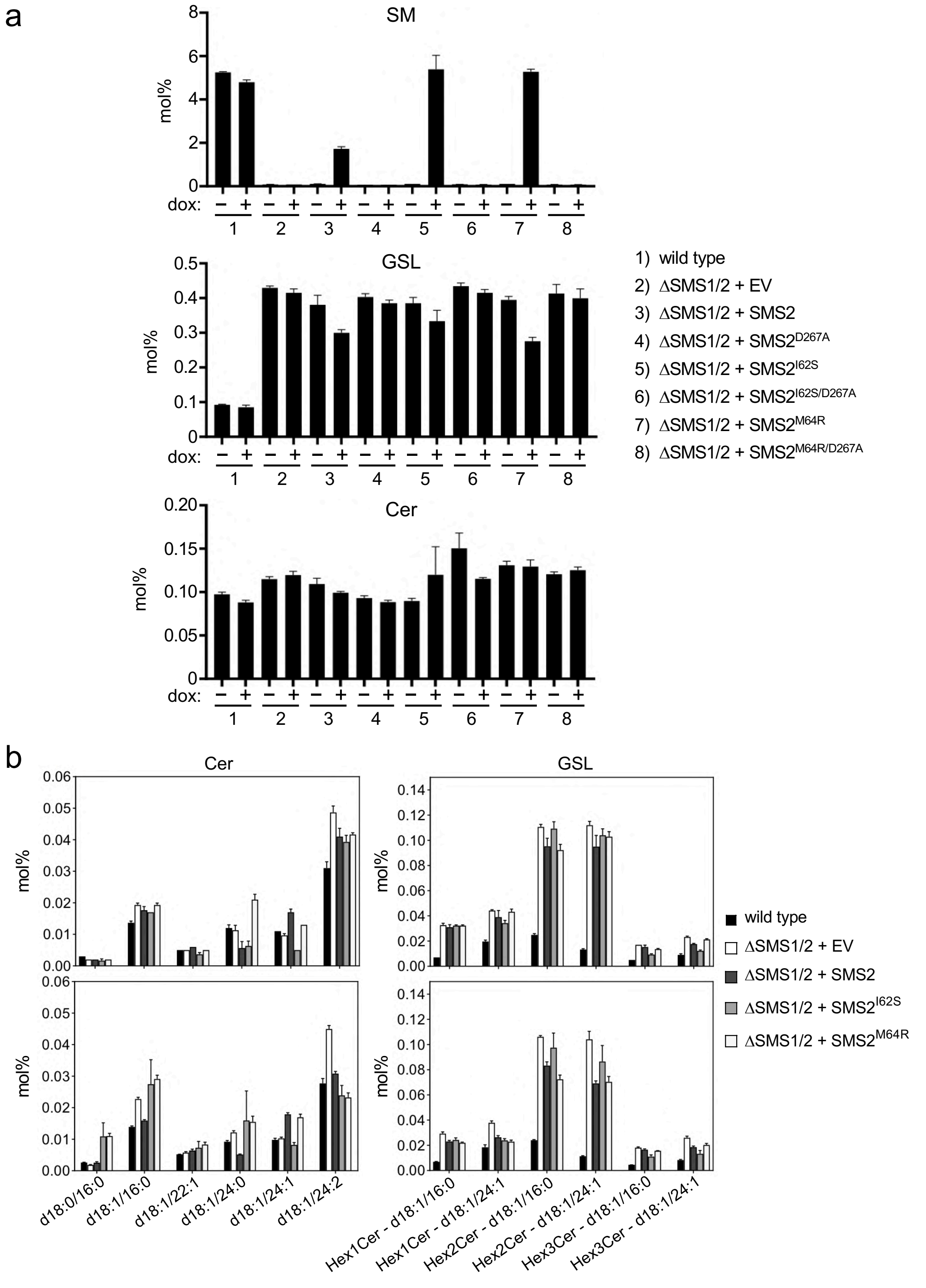


**Supplementary Figure 2. Pathogenic SMS2 variants support bulk production of SM in the ER.** (**a**) HeLa ΔSMS1/2 cells transduced with doxycycline-inducible FLAG-tagged SMS2, SMS2$^{I62S}$, SMS2$^{M64R}$ or their enzyme-dead isoforms (D276A) were grown for 16 h in the absence or presence of 1 µg/ml doxycycline and subjected to total lipid extraction. Cellular SM, glycosphingolipid (HexCer) and ceramide (Cer) levels were quantified by LC-MS/MS and expressed as mol% of total phospholipid analyzed. (**b**) Ceramide and HexCer species in total lipid extracts of cells treated as in (a) were quantified by LC-MS/MS and expressed as mol% of total phospholipid analyzed.

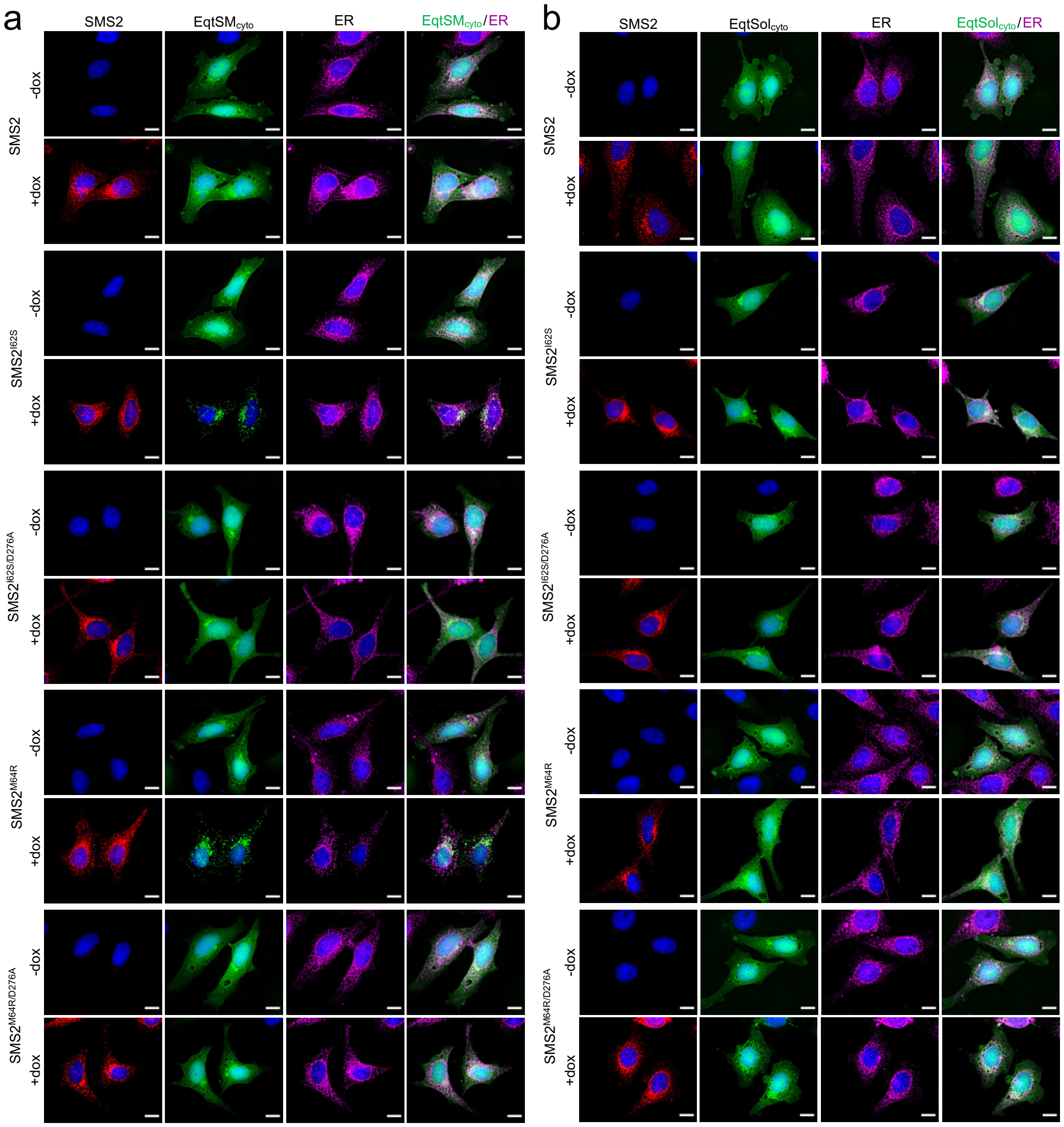


**Supplementary Figure 3. Mobilization of EqtSM$_{cyto}$ by pathogenic SMS2 variants relies on a catalytically active enzyme.**

(**a**) HeLa ΔSMS1/2 cells transduced with FLAG-tagged SMS2, SMS2$^{I62S}$, SMS2$^{M64R}$ or their enzyme-dead isoforms (D276A) were transfected with GFP-tagged EqtSM$_{cyto}$ (*green*) and grown for 16 h in the absence or presence of 1 μg/ml doxycycline. Next, cells were fixed, immunostained with α-FLAG (SMS2, *red*) and α-calnexin (ER, *magenta*) antibodies, counterstained with DAPI (*blue*) and imaged by DeltaVision microscopy. (**b**) Cells as in (a) were transfected with GFP-tagged EqtSol$_{cyto}$ (*green*) and grown for 16 h in the absence or presence of 1 μg/ml doxycycline. Next, cells were fixed, immunostained with α-FLAG (SMS2, *red*) and α-calnexin (ER, *magenta*) antibodies, counterstained with DAPI (*blue*) and imaged by DeltaVision microscopy. Scale bar, 10 μm.

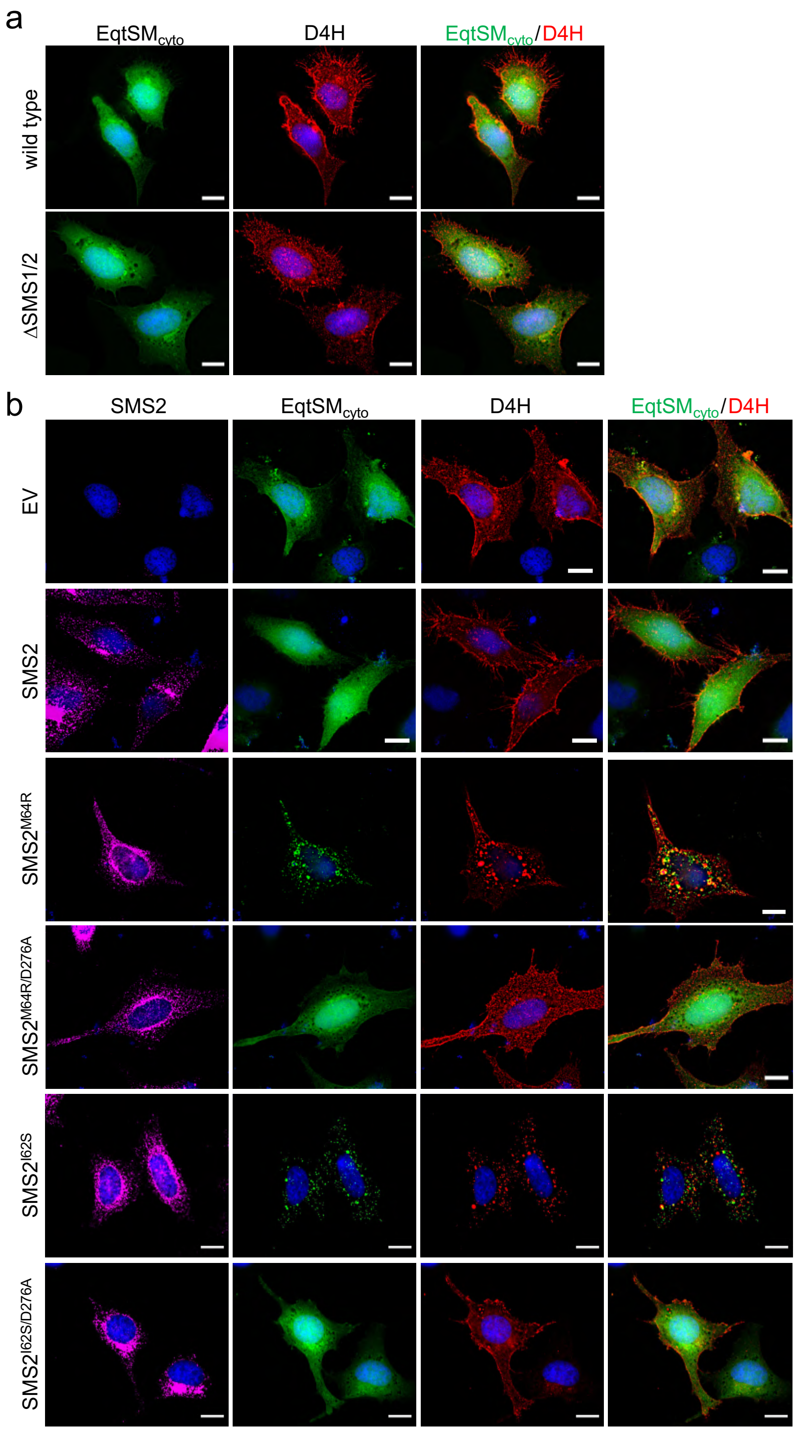


**Supplementary Figure 4. Pathogenic SMS2 variants perturb subcellular cholesterol pools.** (**a**) HeLa wildtype or ΔSMS1/2 cells were co-transfected with GFP-tagged EqtSM$_{cyto}$ (*green*) and mCherry-tagged D4H (*red*). After 16 h, cells were fixed, counterstained with DAPI (*blue*) and imaged by DeltaVision microscopy. (**b**) HeLa ΔSMS1/2 cells transduced with FLAG-tagged SMS2, SMS2$^{I62S}$, SMS2$^{M64R}$ or their enzyme-dead isoforms (D276A) were co-transfected with GFP-tagged EqtSM$_{cyto}$ (*green*) and mCherry-tagged D4H (*red*) and then grown for 16 h in the presence of 1 µg/ml doxycycline. Next, cells were fixed, immunostained with α-FLAG antibodies (SMS2, *magenta*), counterstained with DAPI (*blue*) and imaged by DeltaVision microscopy. Scale bar, 10 µm.

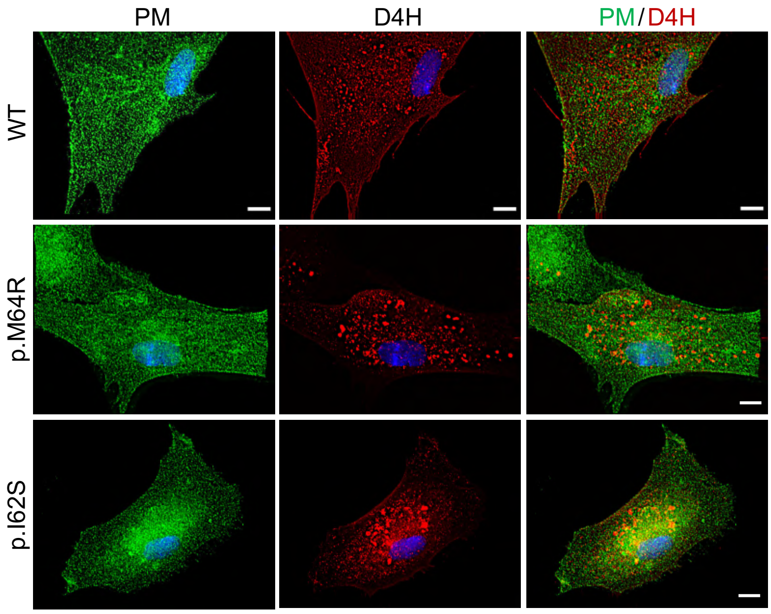


**Supplementary Figure 5. Patient-derived fibroblasts display hogenic SMS2 variants perturb subcellular cholesterol pools.**

Control (WT) or patient-derived human skin fibroblasts carrying heterozygous missense variants c.185T>G (p.I62S) or c.191T>G (p.M64R) in *SGMS2* were transfected with mCherry-tagged D4H (*red*), fixed, immunostained with α-Na/K-ATPase antibodies (PM, *green*) and imaged by Deltavision microscopy. Scale bar, 10 μm.
